# Single-Cell RNA Sequencing Reveals Hemocyte Heterogeneity, Differentiation Trajectories, and Viral Tropism in Shrimp (*Macrobrachium rosenbergii*) Infected with Decapod Iridescent Virus 1

**DOI:** 10.1101/2025.02.10.637476

**Authors:** Bang Xiao, Fang Kang, Taolin Fan, Qianqian Li, ManMan Liu, Junming Pan, Jianguo He, Chaozheng Li

## Abstract

Decapod Iridescent Virus 1 (DIV1) is a highly pathogenic virus threatening crustacean aquaculture. Hemocytes, the primary immune cells in shrimp, play crucial roles in host defense, yet their transcriptional heterogeneity and differentiation dynamics under viral infection remain poorly understood. Here, we characterize hemocyte subpopulations in *Macrobrachium rosenbergii* before and after DIV1 infection at single-cell resolution. We identified 12 distinct hemocyte clusters with unique molecular signatures. DIV1 infection induced significant shifts in hemocyte composition, with clusters 0, 5, 6, and 8 expanding, while clusters 9 and 10 exhibited resistance. Clusters 0, 5, and 8 were highly susceptible, showing elevated viral gene expression. Pseudo-time analysis revealed that DIV1 accelerates hemocyte differentiation, driving prohemocytes and semi-granulocytes toward terminally differentiated granulocytes and hyalinocytes, particularly the immune-active HC4 subpopulation. Functional enrichment analysis showed that susceptible hemocytes were associated with viral processes, oxidative stress, and phagocytosis, while resistant clusters exhibited distinct immune signatures. Furthermore, knockdown experiments confirmed the antiviral roles of *PPO2*, *RAB11B*, *LAMP1*, and *Dorsal*, as their silencing led to increased viral loads, higher MCP protein levels, and reduced shrimp survival. Taken together, our study provides the first single-cell resolution atlas of shrimp hemocytes under DIV1 infection, revealing their transcriptional heterogeneity, differentiation trajectories, and immune responses. These findings offer novel insights into shrimp antiviral immunity and lay the foundation for disease-resistant breeding and immunostimulatory strategies in aquaculture.

**Importances:** Decapod Iridescent Virus 1 (DIV1) is a highly pathogenic virus threatening crustacean aquaculture, causing severe economic losses. Hemocytes, the primary immune cells in shrimp, play crucial roles in antiviral defense, yet their diversity, differentiation, and responses to viral infection remain poorly understood. In this study, we employed single-cell RNA sequencing (scRNA-seq) to comprehensively characterize hemocyte subpopulations in *Macrobrachium rosenbergii* before and after DIV1 infection. We identified 12 transcriptionally distinct hemocyte clusters and revealed significant changes in hemocyte composition following infection. Notably, we discovered that specific clusters (0, 5 and 8) were highly susceptible to DIV1, while others (9 and 10) exhibited resistance. Additionally, we reconstructed hemocyte differentiation trajectories and found that DIV1 infection drives hematopoiesis, accelerating the transition of progenitor cells into terminally differentiated immune-active hemocytes. Functional analysis highlighted key antiviral effectors, including *PPO2*, *RAB11B*, *LAMP1*, and *Dorsal*, which were significantly upregulated in response to infection. These findings provide novel insights into shrimp antiviral immunity and viral pathogenesis, offering potential targets for disease-resistant breeding and immunostimulatory interventions.

## 1. Introduction

Decapod Iridescent Virus 1 (DIV1) is an emerging viral pathogen that has caused significant economic losses in crustacean aquaculture (1–3). DIV1 belongs to the newly established genus *Decapodiridovirus* within the family *Iridoviridae*, a classification approved by the International Committee on Taxonomy of Viruses (ICTV) in 2019. The virus has an icosahedral structure, measuring approximately 140–150 nm in diameter, and possesses a double-stranded DNA (dsDNA) genome of roughly 165 kb (2, 4). DIV1 exhibits a broad host range, infecting both freshwater and marine crustaceans, and induces severe clinical symptoms, including an empty digestive tract, pale hepatopancreas, lethargy, cessation of feeding, and blackened abdominal shell edges (5–7). Infected populations often experience mortality rates approaching 100% (8), highlighting the virus’s high pathogenicity and significant threat to global aquaculture. Given the severe threat posed by DIV1 to aquaculture, a deeper understanding of DIV1-host interactions is crucial for elucidating its pathogenesis and developing effective prevention and treatment strategies. Histopathological studies have demonstrated that hemocytes serve as primary target cells for DIV1 infection (5, 6). In crustaceans, including shrimp, hemocytes play a central role in innate immunity, mediating humoral and cellular defense mechanisms such as antimicrobial peptides (AMPs) production, phagocytosis, encapsulation, nodulation, and apoptosis (9, 10). Traditionally, shrimp hemocytes have been classified into three main types based on morphology and granule staining: hyalinocytes (HC), granulocytes (GC), and semi-granulocytes (SGC) (11–13). However, these classifications remain ambiguous due to the limitations of morphological analysis, leading to discrepancies in their immune functions. Moreover, how DIV1 infection influences hemocyte differentiation and functional specialization remains unclear. Understanding the relationship between viral infection and hemocyte differentiation is crucial for unraveling host immune strategies and viral pathogenesis.

Although previous omics studies have explored the molecular functions of shrimp hemocyte subpopulations, their low resolution has limited the identification of rare and low-abundance cell types (7, 8, 14). Consequently, the heterogeneity of hemocyte immune responses and the developmental trajectories of these immune cells remain poorly understood. Furthermore, previous studies have not determined which hemocyte subpopulations are most susceptible to DIV1 infection and which exhibit resistance. Investigating the viral tropism among hemocyte clusters will provide key insights into host-virus interactions and potential mechanisms of immune evasion.

Single-cell RNA sequencing (scRNA-seq) has emerged as a powerful tool for characterizing dynamic gene expression patterns at the single-cell level. Unlike bulk RNA-seq, which averages gene expression across heterogeneous cell populations, scRNA-seq enables the precise identification of hemocyte subtypes, the discovery of rare immune cell populations, and the reconstruction of differentiation trajectories (15–17). ScRNA-seq has been extensively used in vertebrates to elucidate cellular responses to viral infections, such as SARS-CoV-2 (18–20). In aquatic species, scRNA-seq has been applied to investigate immune responses in fish infected with red grouper neuron necrosis virus (RGNNV), revealing neuronal targets such as GLU1 and GLU3 (21). Moreover, scRNA-seq has been successfully utilized to classify hemocyte populations in invertebrates, including *Drosophila*, mosquitoes, oysters, and shrimp (22–25). Beyond immune profiling, scRNA-seq has proven effective in identifying viral target cells by detecting viral mRNA within host cell populations (26–28). For instance, scRNA-seq has been used to map the immune response landscape in silkworm larvae following *Bombyx mori* nucleopolyhedrovirus (BmNPV) infection (29). These studies demonstrate the unique advantages of scRNA-seq in resolving cellular heterogeneity and host-virus interactions.

In this study, we employed scRNA-seq to comprehensively characterize the heterogeneity of hemocytes in *Macrobrachium rosenbergii*, a natural and susceptible host of DIV1, and to examine their transcriptional responses to infection. Our analysis identified 12 distinct hemocyte clusters, each with unique transcriptional signatures and functional roles. We further uncovered differential viral tropism among these clusters, with clusters 0, 5 and 8 being highly susceptible to DIV1, while clusters 9 and 10 exhibited resistance. Key immune genes, including *PPO2*, *RAB11B*, *LAMP1*, and *Dorsal*, were found to play critical roles in antiviral defense. These findings provide novel insights into the cellular and molecular mechanisms underlying shrimp immunity and host-virus interactions.

## 2. Results

### 2.1 Identification of Hemocyte Clusters in *M. rosenbergii* Using scRNA-seq

To investigate the functional subtypes, differentiation characteristics, and immune responses of hemocytes in *M. rosenbergii* to DIV1 infection, hemocytes were collected at 0-, 4-and 24-hours post-infection (hpi) and subjected to single-cell RNA sequencing (scRNA-seq). After filtering low-quality data during preprocessing, a total of 48,042 high-confidence cells were obtained from the sequencing data (Genome Sequence Archive, PRJCA035213) (Fig. 1A). The distribution of cell counts across the six groups was as follows: 5,840 (CK-1), 8,787 (CK-2), 8,086 (DIV1-4h-1), 8,835 (DIV1-4h-2), 7,835 (DIV1-24h-1), and 8,659 (DIV1-24h-2). The total number of unique molecular identifiers (UMIs) captured in each sample was 1.41 × 10⁷ (CK-1), 2.42 × 10⁷ (CK-2), 2.50 × 10⁷ (DIV1-4h-1), 2.49 × 10⁷ (DIV1-4h-2), 2.88 × 10⁷ (DIV1-24h-1), and 3.40 × 10⁷ (DIV1-24h-2), respectively (Supplementary Table 2).

**Figure 1.**
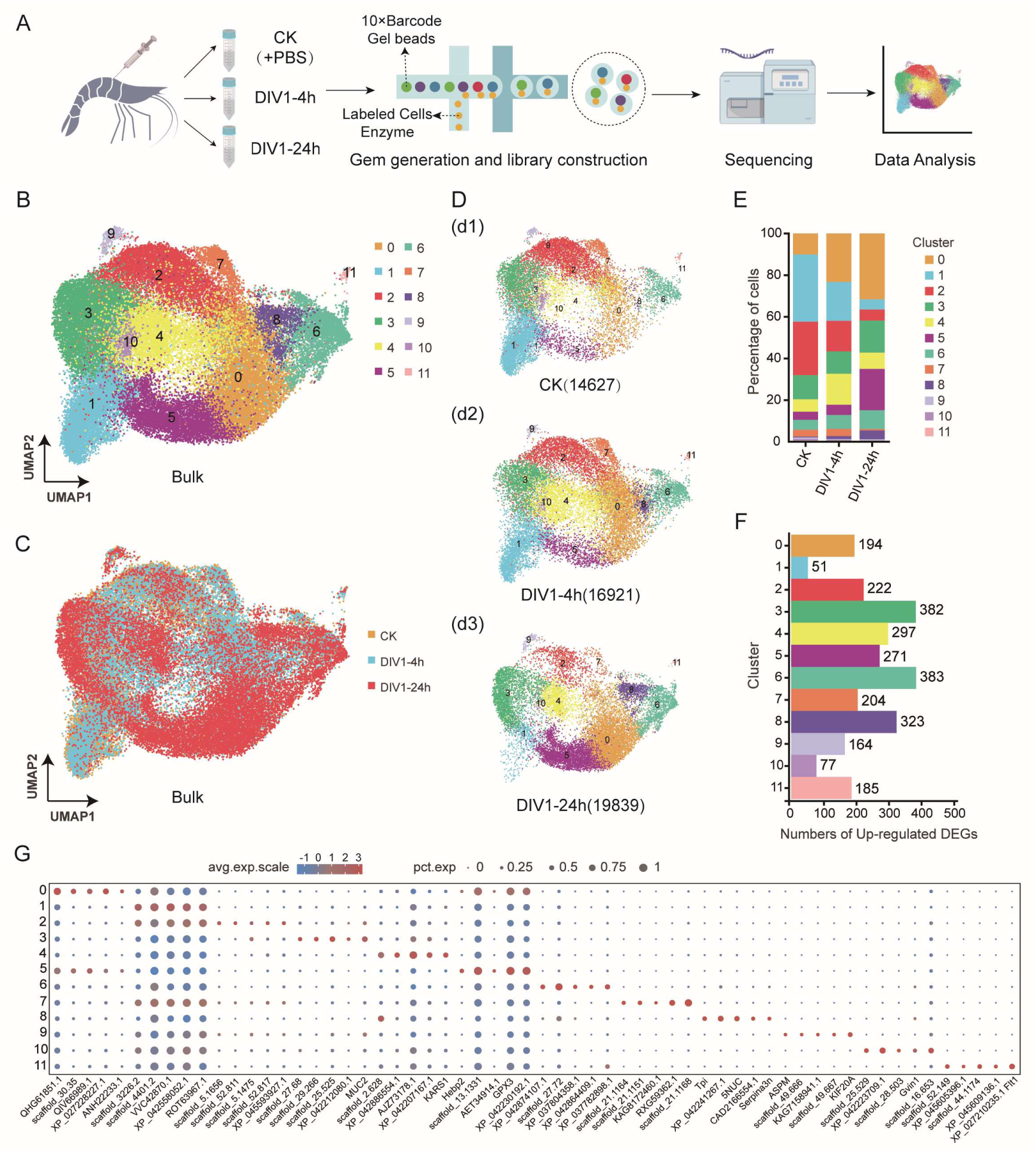
Single cell profiling of hemocytes collected from DIV1-infected and healthy shrimp *M. rosenbergii*. (A) The schematic of scRNA-Seq and 10 × genomic data analyses (Genome Sequence Archive, PRJCA035213). (B) UMAP projection representing the 12 cell clusters identified in the shrimp hemocytes (unified set of control and DIV1 infection samples). (C) UMAP plots representing clusters of hemocyte from pre-DIV1 infection (yellow) to post-DIV1 infection (blue for early stage and red for late stage of infection). (D) UMAP plots representing the cell profiles of CK group (d1), DIV1-4h group (d2) and DIV1-24h group (d3), respectively. (E) Proportion of cells in each cluster and their distribution in the healthy and DIV1-infected samples. (F) The number of differential expression genes (DEGs) in each cluster. (G) Dot plot illustrating the top five DEGs enriched in each cluster, based on avg. logFC. The size of each dot indicates the percentage of cells expressing any gene within the respective cluster. The color gradient of the dots reflects the expression level relative to other clusters.

Next, using Uniform Manifold Approximation and Projection (UMAP) for dimensionality reduction and unsupervised clustering, we identified 12 distinct hemocyte clusters (named clusters 0∼11) with unique gene expression profiles based on the scRNA-seq data (Fig. 1B). The UMAP visualization highlighted clear differences in the distribution of hemocyte clusters between CK, DIV1-4h, and DIV1-24h groups (Figs. 1C, 1D). Analysis of cluster composition revealed significant changes in the proportions of specific hemocyte clusters following DIV1 infection (Fig. 1E). Notably, the proportions of clusters 0, 5, 6, and 8 increased progressively with the duration of DIV1 infection, while clusters 1, 2, 7, and 10 showed marked decreases at 24 hpi (Fig. 1E). Additionally, the number of upregulated genes detected in each hemocyte cluster varied significantly. Clusters 3, 6, and 8 exhibited the highest numbers of upregulated genes, suggesting their active involvement in immune responses (Fig. 1F). To further explore the roles of these clusters, the top five differentially expressed genes (DEGs) in each hemocyte cluster were identified and their expression abundances visualized in a dot plot (Fig. 1G; Supplementary Table 3). Collectively, these results demonstrate the functional heterogeneity of hemocytes in *M. rosenbergii* and their distinct immune functions during DIV1 infection.

### 2.2 Functional Enrichment Analysis of Hemocyte Clusters

To further elucidate the biological roles of each hemocyte cluster, functional enrichment analyses of differentially expressed genes (DEGs) were performed using Gene Ontology (GO) and KEGG databases (Supplementary Tables 4, 5). GO terms such as “immune system process,” “response to stimulus,” and “viral process” were significantly enriched in clusters 3, 4, 5, 6, 8, and 11 compared to other clusters. These findings were consistent with KEGG pathway annotations (Fig. 2A). Pathways related to immune responses, including “Immune System,” “Infectious Diseases: Viral,” and “Signal Transduction,” were primarily enriched in clusters 3, 6 and 8 (Fig. 2B).

**Figure 2.**
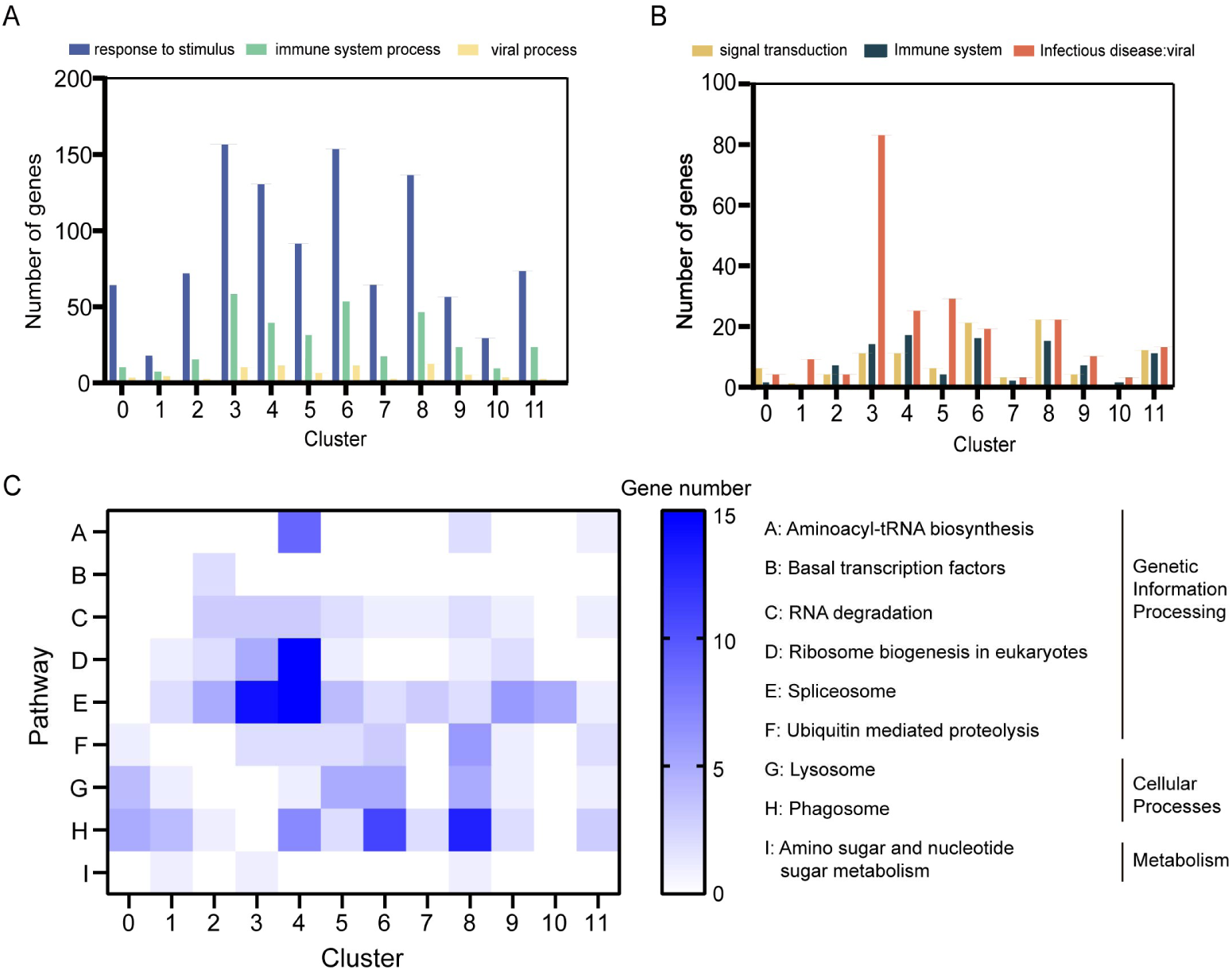
Plot profiling of the GO analysis and KEGG pathway enrichment analysis in each cluster. (A) The numbers of genes involved in the “immune system process”, “response to stimulus”, and “viral process” of GO terms as estimated in each cluster. (B) The numbers of genes involved in the “Immune System,” “Infectious Diseases: viral” and “Signal Transduction” pathways in KEGG as analyzed in each cluster. (C) The enrichment of genes in each cell cluster in pathways related to regulating of gene transcription and expression, phagocytosis and carbohydrate metabolism.

DEGs associated with metabolic pathways, including lipid, amino acid, and glucose metabolism, were predominantly enriched in clusters 3 and 5 (Supplementary Table 5). Additionally, genes involved in gene duplication and repair processes were found in clusters 0, 3, 5, 8, 10, and 11, indicating the presence of proliferating cells in these clusters (Supplementary Table 5). Genes related to cell growth and death processes were distributed across all clusters but were especially enriched in clusters 3, 4, 6, and 8, highlighting their role in regulating cell development under DIV1 infection (Supplementary Table 5).

KEGG pathway analysis further revealed that transcription and translation regulation pathways, including “ribosome biogenesis in eukaryotes,” and “spliceosome” were significantly enriched in clusters 3 and 4 compared to other clusters (Fig. 2C). Differently, lysosomal and phagosomal pathways were highly enriched in clusters 6 and 8 (Fig. 2C), suggesting their involvement in immune responses, particularly in phagocytosis. Interestingly, pathways related to carbohydrate metabolism, such as aminoglycan metabolism and ribose metabolism was not significantly enriched in any cluster, reflecting a potentially limited role of these pathways during DIV1 infection (Fig. 2C).

### 2.3 Functional Characterization of Hemocyte Clusters During DIV1 Infection

To gain deeper insights into the immune functions of distinct hemocyte subpopulations during DIV1 infection, we analyzed differentially upregulated genes (DEGs) across the 12 identified hemocyte clusters. Based on their gene expression profiles and functional annotations, these clusters could play distinct roles in the immune response against DIV1.

Clusters 2, 7, and 9 were enriched with genes related to cell proliferation, division, and germline development, including *My018a*, *DOCK3*, *EIF4ENIF1*, *AKAP9*, *ASPM*, *INCENP*, and *KIF20A* (Supplementary Fig. 1A). Among these, *ASPM* is a known marker of granulocyte-monocytic progenitor cells in humans (30), while *INCENP* and *KIF20A* are highly expressed in basal cells of mouse tracheal circulation (31). This suggests that these clusters may represent progenitor-like or proliferative hemocytes involved in cell renewal.

Clusters 3, 4, and 10 exhibited high expression of genes associated with cytoskeletal formation, immune stress, and cell development, such as *MUC2*, *SOD2*, *BYSL*, *Hspe1*, *HSP90AB1*, *DIRAS2*, *GCSH*, *tun*, *CDSP2L*, *ced-10*, *SPE*, *Rel*, and *Casp2* (Supplementary Figs. 1B, 1C). In cluster 3, *MUC2* and *SOD2* were linked to cytoskeletal organization, affecting cell differentiation, migration, and aggregation (32, 33). Additionally, lysosome-related genes such as *Tun* and *GCSH* were highly expressed in cluster 3, suggesting its potential for phagocytic activity. Cluster 4 was enriched in key crustacean innate immunity pathways, including *Toll*, *IMD*, and *JAK/STAT*, which play vital roles in responding to pathogen invasion (34). Although cluster 10 shared functional similarities with cluster 4, it lacked a highly specific marker gene (Supplementary Fig. 1C).

Clusters 1 and 11 were characterized by high expression of ribosomal protein-related genes, such as *RPS3*, *RPS7*, and *RpL10*, along with translation initiation factor genes, including *eif3eb* and *v1g240395* (Supplementary Fig. 1D). These clusters were associated with enhanced protein synthesis in response to viral infection. Notably, *CD63*, a marker highly expressed in macrophages and monocytes in other species (33), was also upregulated in cluster 11 (Supplementary Fig. 1D).

Clusters 0, 5, 6, and 8 were closely linked to immune signaling, the prophenoloxidase (proPO) system, and antimicrobial peptides (AMPs) production. The proportions of cells in these clusters increased steadily after DIV1 infection (Fig. 3A; Supplementary Table 2), suggesting their significant role in antiviral defense. In clusters 0 and 5, key immune-related genes such as *PPAF*, *Crustin*, and *GPX3* were highly enriched (Fig. 3B), highlighting their involvement in innate immunity and viral elimination (35). In clusters 6 and 8, genes homologous to specific markers of mouse macrophages (*FCN1*, *Tie2*) (36) and natural killer T cells (*CACT*, *TPI*) in humans (37) were highly expressed (Fig. 3C).

**Figure 3.**
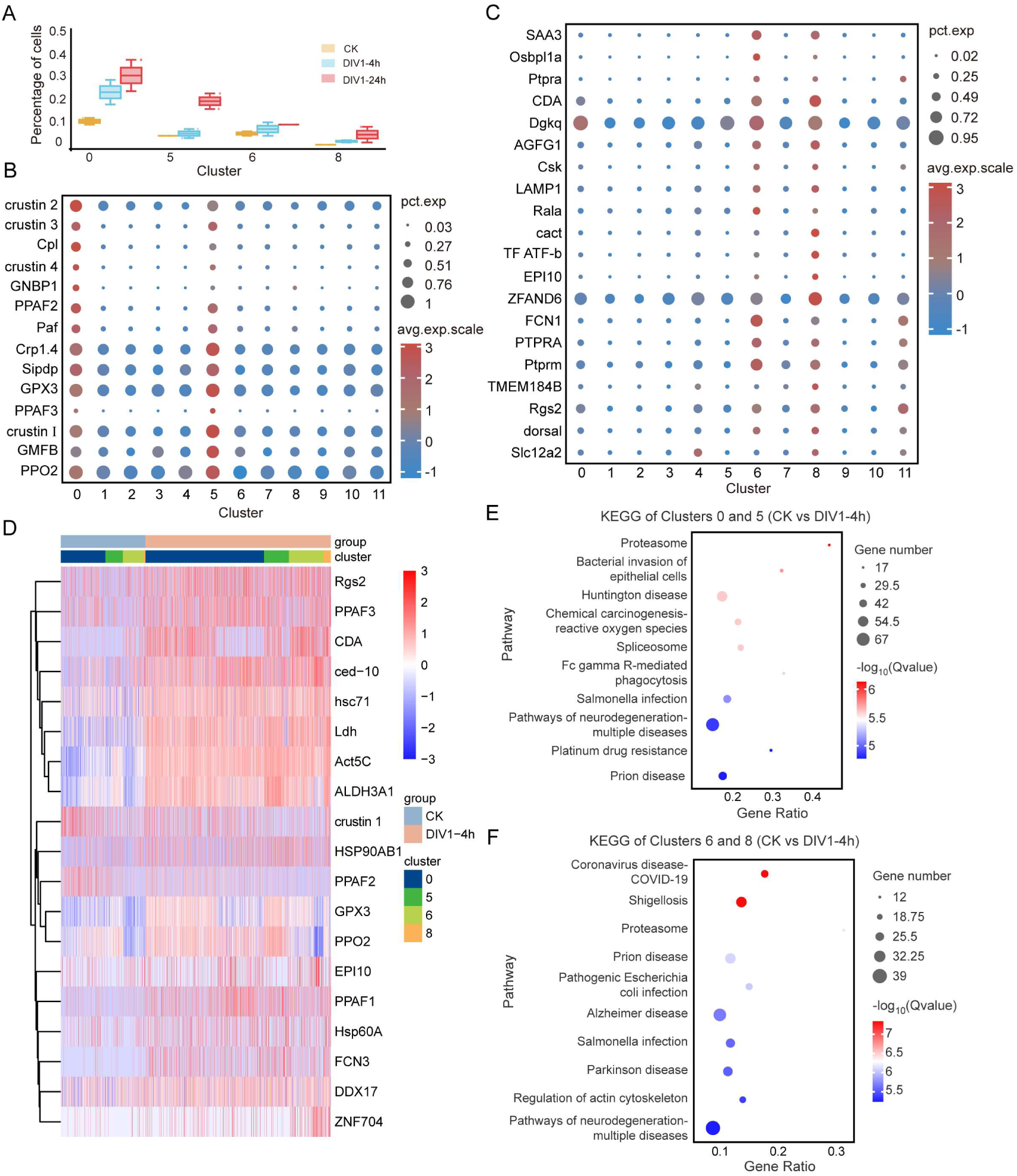
Molecular characterization of the clusters 0, 5, 6, and 8. (A) The cell proportions of 0, 5, 6 and 8 cell clusters were continuously upregulated across three experimental conditions: CK (control), DIV1-4h, and DIV1-24h. (B) Dot plot showing the top immune genes enriched in clusters 0 and 5. The dot size represents the percentage of cells expressing the gene, and the color indicates the average scaled expression level. (C) Dot plot representing the top immune genes enriched in clusters 6 and 8. The dot size represents the percentage of cells expressing the gene, and the color indicates the average scaled expression level. (D) Heatmap of representative gene expression patterns in clusters 0, 4, 5, and 8, compared to the control group after DIV1 infection at 4h. (E, F) TOP 10 KEGG pathway enrichment analysis of differential genes in clusters 0 and 5 (E) /clusters 6 and 8 (F) compared to the control group at 4 hpi.

Further analysis of DEGs between the control and DIV1-infected groups showed upregulated expression of *PPAF1*, *PPAF2*, *PPAF3*, and *PPO2* in clusters 0 and 5 (Fig. 3D). In clusters 6 and 8, zinc finger protein *ZNF704* was significantly upregulated (Fig. 3D; Supplementary Fig. 2A), which is known to degrade viral RNA or assist in folding viral proteins (38). Apoptosis-related genes such as *ced-10* and heat shock proteins (*hsc71*, *Hsp60A*, *HSP90AB1*) were also upregulated in clusters 0 and 6 (Fig. 3D; Supplementary Fig. 2B), indicating their involvement in programmed cell death during viral infection (39, 40). KEGG analysis confirmed that clusters 0, 5, 6, and 8 were pivotal in immune defense during the early and late stages of DIV1 infection (Figs. 3E, 3F; Supplementary Figs. 1E, 1F).

Moreover, comparative analysis revealed that distinct cell clusters exhibited a substantial number of significantly upregulated genes across multiple stages following DIV1 infection when contrasted with the healthy control group (Supplementary Figs. 3A, 3B). Notably, pronounced molecular heterogeneity was observed between later stages versus early stages of DIV1 infection within these cellular clusters (Supplementary Fig. 3C), demonstrating temporal progression of transcriptional reprogramming.

Together, these findings demonstrate that different hemocyte clusters play distinct and complementary roles in combating DIV1 infection. Clusters 0, 5, 6, and 8 were identified as the primary contributors to antiviral immunity, while other clusters supported processes such as cell renewal and protein synthesis.

### 2.4 Differential Viral Tropism of Hemocyte Clusters to DIV1

To characterize the viral tropism of different hemocyte clusters, the scRNA-seq data were mapped to the DIV1 genome. The UMAP plots illustrated the cell atlas containing 12 clusters of shrimp hemocytes and transcriptional profiles of DIV1 genes across the hemocyte clusters (Supplementary Fig. 4A), revealing distinct patterns of viral gene expression between clusters at 24 hpi, despite the relatively low viral load during the early stages of infection (Supplementary Fig. 4B; Supplementary Table 6). We found that clusters 9 and 10 did not express any detectable DIV1 genes, suggesting that these clusters could be resistant or insensitive to DIV1 infection (Figs. 4A, 4B). In contrast, varying levels of viral gene expression were observed in the remaining hemocyte clusters, with clusters 0 and 5 exhibiting the highest transcriptional levels of DIV1 genes. This indicates that clusters 0 and 5 are the most susceptible to DIV1 infection (Figs. 4A, 4B). Heatmap analysis of the top 42 highly expressed viral genes across clusters supported these findings, highlighting the preferential susceptibility of clusters 0 and 5 (Fig. 4C).

**Figure 4.**
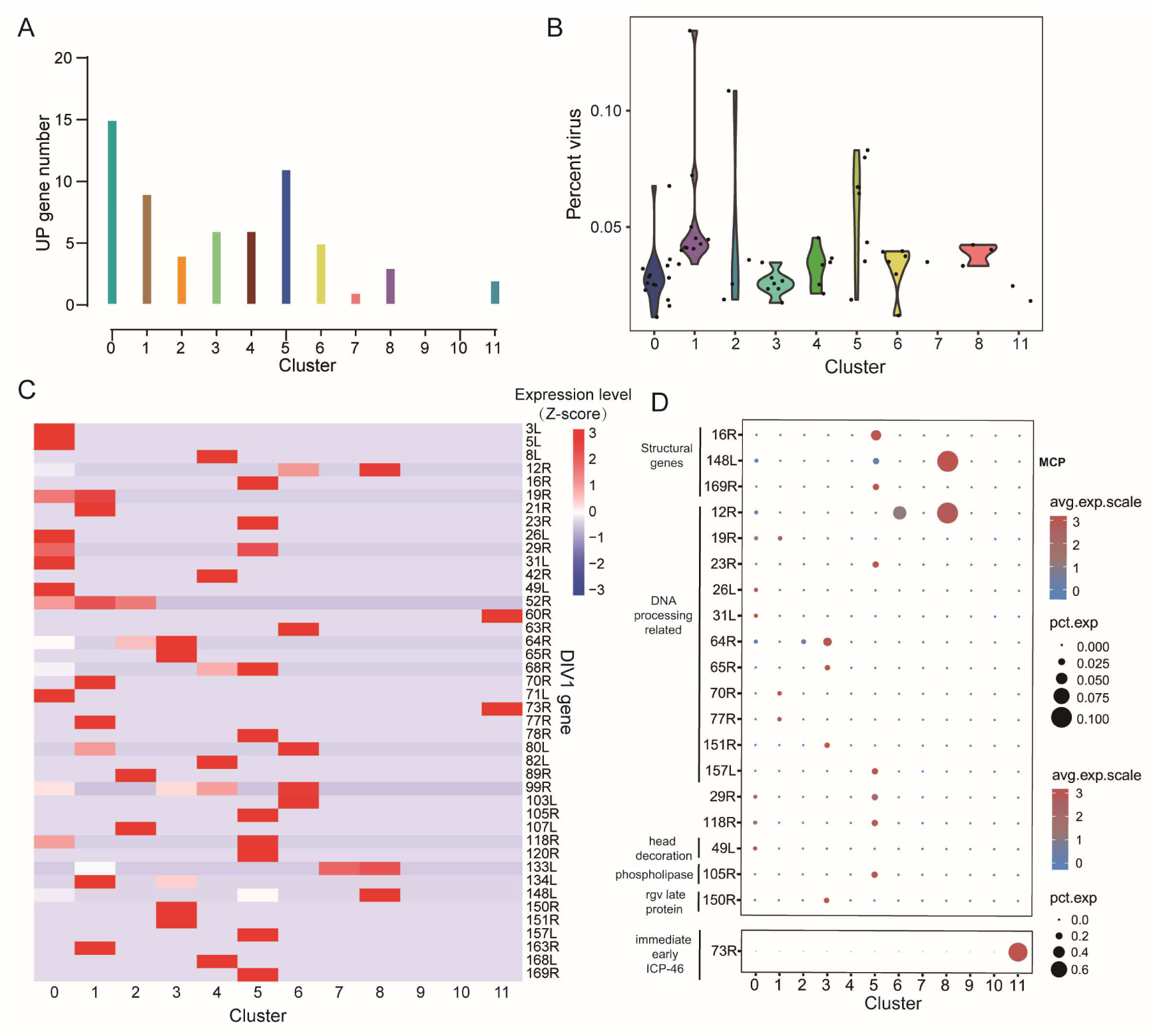
Analysis of viral gene expression and viral load in hemocyte cluster post-DIV1 infection. (A) Statistics on the number of specific viral genes in each cluster. (B) Violin plots showing the cells with different virus load (UMI_DIV1_/(UMI_DIV1_ + UMI_host_)) in each cluster except clusters 9, 10 which do not have any virus genes captured. (C) Heatmap showing the normalized expression (Z-score) of all expressed DIV1 genes in each cell cluster of the DIV1-infected groups. (D) Bubble plot showing the expression profile of key virus genes in each cluster.

To further elucidate the infection dynamics, viral genes were categorized into three temporal phases—immediate early (IE), early, and late—based on their expression patterns during the viral life cycle. Among these, the DIV1-IE gene ICP-46 (*73R*) was highly expressed in cluster 11, suggesting that cells in this cluster are more prone to initiating viral replication (Fig. 4D). In clusters 5 and 8, late-stage viral structural protein genes such as *169R*, *16R*, and *148L* were highly expressed (Fig. 4D). Notably, *169R* encodes a transmembrane envelope protein implicated in mediating host-virus interactions (41), while *148L* encodes the major capsid protein (MCP), a conserved structural protein essential for viral assembly (42).

Together, these findings reveal that hemocyte clusters exhibit varying levels of susceptibility to DIV1 infection. While clusters 0, 5, and 8 were highly susceptible, clusters 9 and 10 appeared resistant. The expression patterns of viral genes across clusters highlight the diverse roles of hemocytes in the host-virus interaction and suggest potential mechanisms underlying cluster-specific susceptibility.

### 2.5 Expression Profiles of Immune Genes in Clusters 0, 5, 6, and 8 During DIV1 Infection

To investigate the immune responses of clusters 0, 5, 6, and 8, which were identified as the primary immune-active hemocyte clusters during DIV1 infection, we analyzed the differentially upregulated genes (DEGs) across these clusters at 0, 4 and 24 hpi using STEM analysis. And we prioritized profiles 5, 6, and 7 among the eight transcriptional patterns, as these profiles demonstrated significant enrichment of genes associated with active antiviral response pathways (Fig. 5A; Supplementary Fig. 5). A cross-comparison of DEGs in clusters 0 and 5 revealed a consistent upregulation of genes such as *RAB11B*, *PPO2*, and *GPX3*, which displayed a sustained increase in expression across both time points (Figs. 5B, 5C, 5D). Among these, *RAB11B* was notably upregulated not only in clusters 0 and 5 but also in other clusters, suggesting its broad involvement in the immune response to DIV1 (Supplementary Table 7). In clusters 6 and 8, immune-related genes such as *SAA3*, *Dorsal*, and *LAMP1* were highly expressed (Figs. 5B, 5C, 5D). Additionally, *14-3-3ζ* and *ITGA* were specifically and consistently upregulated in cluster 8, while their upregulation in clusters 0 and 5 was restricted to the early stages of infection (Fig. 5B). These results highlight the distinct and time-dependent expression profiles of immune genes in these clusters.

**Figure 5.**
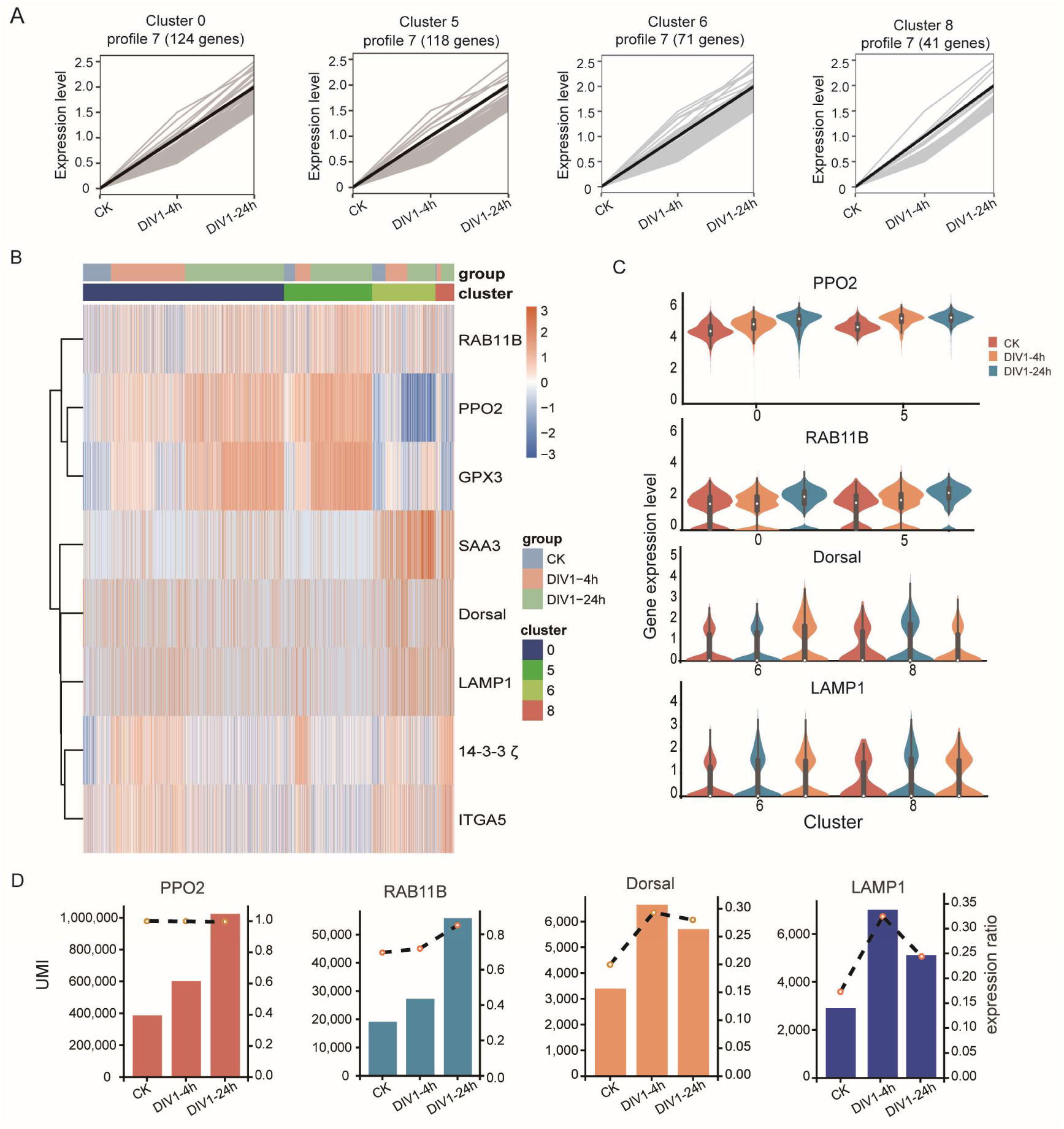
The expression profiles of immune genes in clusters 0, 5, 6, and 8 in response to DIV1 infection. (A) Line plots showing the expression profiles of genes across different time points (CK, DIV1-4h, and DIV1-24h) in cluster 0, cluster 5, cluster 6, and cluster 8. The black lines represent the average expression trend within each cluster, while gray lines depict individual gene trajectories. (B) Heatmap representing the expression levels of key marker genes, across different clusters (color-coded) and experimental groups (CK, DIV1-4h, DIV1-24h) in clusters 0, 5, 6, and 8. Higher expression levels are indicated in red, and lower levels in blue. (C) Violin plots of selected marker genes across conditions and clusters: *MrPPO2, MrRAB11B, MrDorsal*, and *MrLAMP1*. (D) The expression of *MrPPO2, MrRAB11B, MrDorsal*, and *MrLAMP1* across experimental groups. The column height indicates the UMI value detected by the genes, and the dotted line represents the relative expression abundance of the genes.

To validate the functional roles of key DEGs, we performed *in vivo* knockdown experiments for *PPO2*, *RAB11B*, *LAMP1*, and *Dorsal* during DIV1 infection. For *PPO2* knockdown (Fig. 6A), shrimp with silenced *PPO2* exhibited a significantly higher viral load in hemocytes (Fig. 6B) and elevated protein levels of the major capsid protein MCP (Fig. 6C). Additionally, the survival rate of *PPO2*-silenced shrimp infected with DIV1 was significantly lower than that of the control group (Fig. 6D). For *RAB11B* knockdown (Fig. 6E), *RAB11B*-silenced shrimp displayed an increased viral load in hemocytes (Fig. 6F) and elevated MCP protein levels (Fig. 6G). The survival rate of *RAB11B*-silenced shrimp infected with DIV1 was also significantly reduced compared to the control group (Fig. 6H). For *LAMP1* knockdown (Fig. 6I), *LAMP1*-silenced shrimp exhibited a higher viral load in hemocytes (Fig. 6J) and increased MCP protein levels (Fig. 6K). Correspondingly, their survival rate was significantly lower than that of the control group (Fig. 6L). For *Dorsal* knockdown (Fig. 6M), shrimp with silenced *Dorsal* showed a greater viral load in hemocytes (Fig. 6N) and increased MCP protein levels (Fig. 6O), along with a significantly reduced survival rate compared to the control group (Fig. 6P). Moreover, knockdown of *PPO2*, *RAB11B*, *LAMP1*, and *Dorsal* resulted in more severe hepatopancreatic damage post-DIV1 infection compared to the control group (Fig. 6Q). These findings demonstrate that *PPO2*, *RAB11B*, *LAMP1*, and *Dorsal* are critical immune effectors in the response to DIV1 infection. Their upregulation in specific hemocyte clusters underlines their essential roles in restricting viral replication and mitigating host tissue damage.

**Figure 6.**
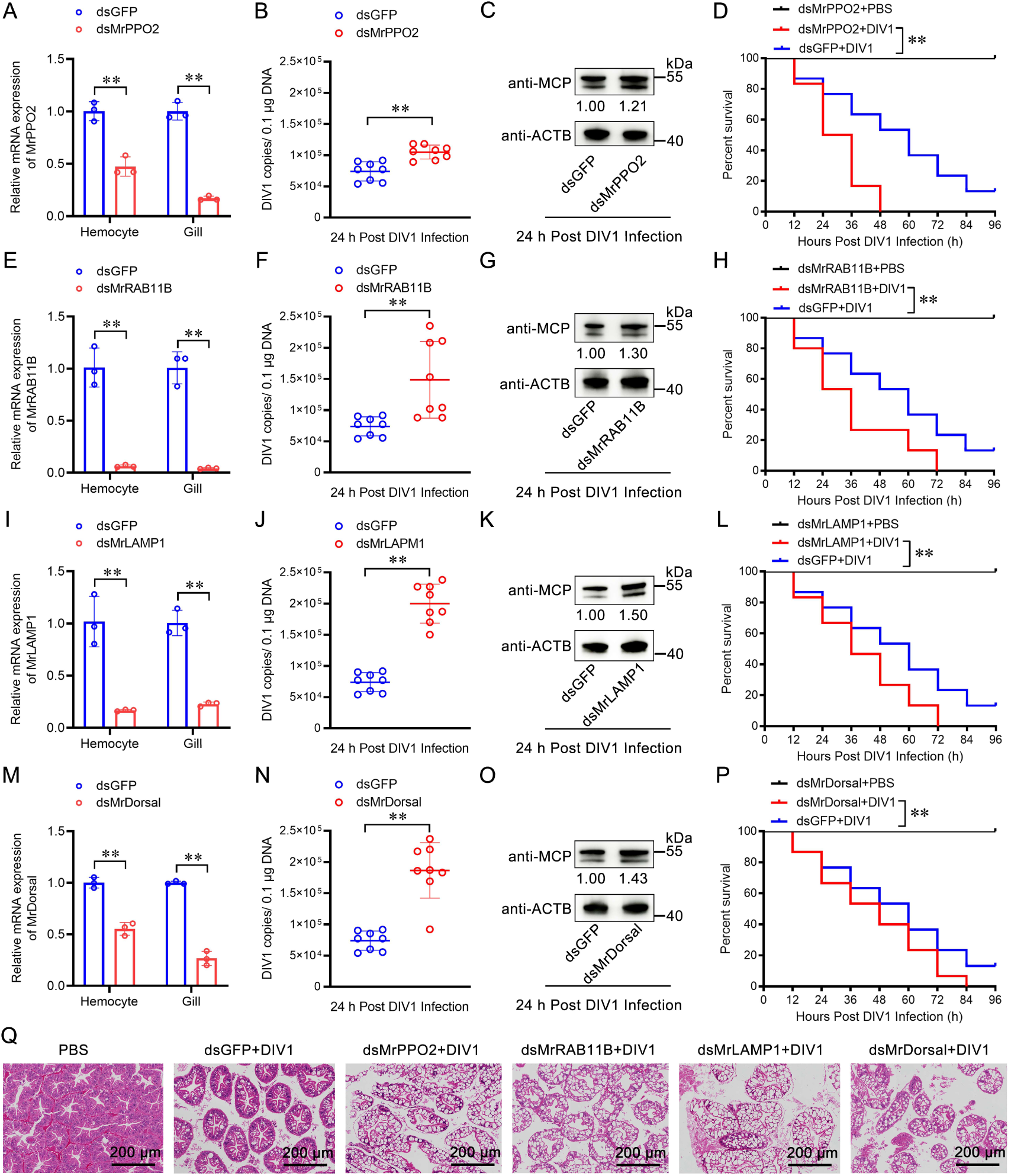
Role of the immune genes in restricting DIV1 replication. (A-D) Role of *MrPPO2* in restricting DIV1 replication. The silencing efficiency of *MrPPO2* (A). The quantity of DIV1 copies was determined by absolute quantitative PCR (B). DIV1 MCP protein was detected by Western blotting (C). The survival rates of DIV1-infected shrimp post-*MrPPO2* knockdown (D). (E-H) Role of *MrRAB11B* in restricting DIV1 replication. The silencing efficiency of *MrRAB11B* (E). The quantity of DIV1 copies was determined by absolute quantitative PCR (F). DIV1 MCP protein was detected by Western blotting (G). The survival rates of DIV1-infected shrimp post-*MrRAB11B* knockdown (H). (I-L) Role of *MrLAMP1* in restricting DIV1 replication. The silencing efficiency of *MrLAMP1* (I). The quantity of DIV1 copies was determined by absolute quantitative PCR (J). DIV1 MCP protein was detected by Western blotting (K). The survival rates of DIV1-infected shrimp post-*MrLAMP1* knockdown (L). (M-P) Role of *MrDorsal* in restricting DIV1 replication. The silencing efficiency of *MrDorsal* (M). The quantity of DIV1 copies was determined by absolute quantitative PCR (N). DIV1 MCP protein was detected by Western blotting (O). The survival rates of DIV1-infected shrimp post-*MrDorsal* knockdown (P). (Q) Morphology analysis of the hepatopancreases with DIV1 infection post the immune genes knockdown. Scale bar, 200 μm. Statistical analysis was performed using the Kaplan-Meier plot (log-rank χ^2^ test). All experiments were conducted with three biological replicates, consistently yielding similar results.

### 2.6 Functional Differentiation of Hemocytes During DIV1 Infection

To explore how DIV1 infection impacts hemocyte differentiation, we constructed cell differentiation trajectories using Monocle 2. The analysis divided the differentiation trajectory into three main branches: a lineage comprising semi-granular cells (SGC) and prohemocytes (PH; yellow circle), a lineage comprising granular cells (GC) and hyalinocytes (HC; purple circle), and a separate lineage predominantly of GC (red circle) (Fig. 7A). Pseudotime analysis demonstrated that distinct hemocyte clusters occupied specific positions along these differentiation trajectories, reflecting significant differences in their differentiation status between infected and uninfected groups (Figs. 7A, 7B).

**Figure 7.**
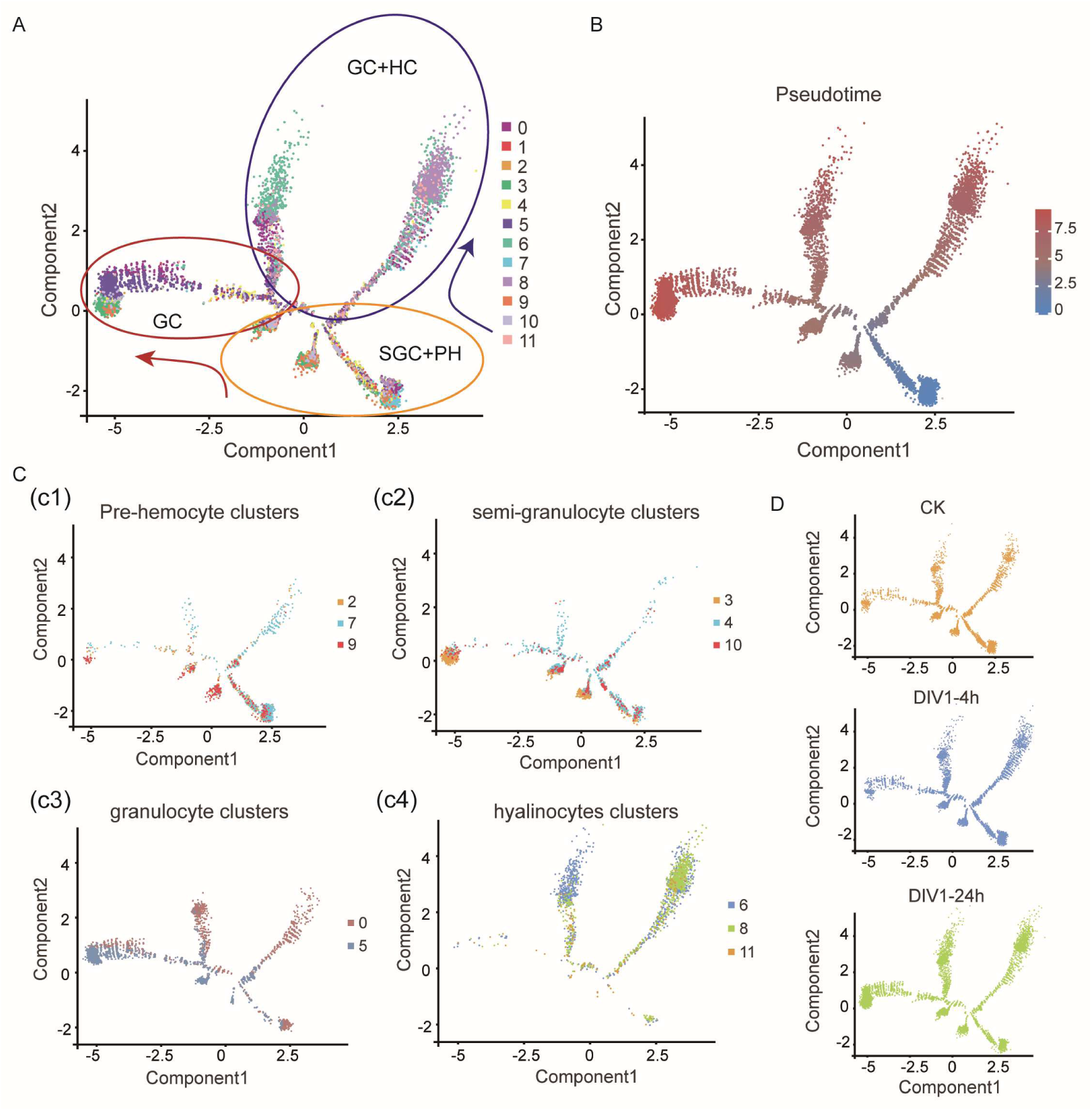
Pseudo-time trajectories of hemocyte clusters. (A) The differentiation trajectory of each cell cluster was constructed using Monocle2. grouped into broader categories: germinal center cells (GC), germinal center plus hemocytes (GC+HC), and semi-granulocyte clusters plus pro-hemocytes (SGC+PH). The arrows highlight directional transitions between categories. (B) Pseudo-time trajectory analysis of all clusters, as the color changes from blue to red, the value of the pseudo-time increases. (C) UMAP plots displaying the major functional groups of clusters: pre-hemocyte clusters (2, 7, 9) (c1), semi-granulocyte clusters (3, 4, 10) (c2), granulocyte clusters (0, 5) (c3), and hyalinocyte clusters (6, 8, 11) (c4). (D) UMAP plots showing temporal dynamics of cluster distribution under DIV1 infection: control (CK), DIV1-4h, and DIV1-24h.

Next, the trajectory analysis revealed four main branches. Cells in the SGC and PH lineage exhibited highly overlapping distribution patterns, while GC and HC lineages showed partial overlap, suggesting the presence of transitional cells co-expressing GC and HC marker genes (Fig. 7C). Importantly, the number of cells in the GC and HC branches increased significantly following DIV1 infection, consistent with earlier findings that DIV1-induced stress promotes the differentiation of SGC and PH into terminally differentiated cells (Fig. 7D). Expression profiles of key differentiation-related genes, including *LAMP1* and transcription factors such as *TAF3* and *Fib*, showed an upregulation trend in the first and second lineages (HC+GC lineage1/2, corresponding to differentiation States 1 and 7). This was consistent with the high expression of *LAMP1* in clusters 6 and 8 (Supplementary Figs. 6A–6E). These results indicate that DIV1 infection significantly alters hemocyte differentiation, driving SGC and PH cells to transition into GC and HC cells, with distinct transcriptional dynamics.

### 2.7 Re-clustering and Immunological Characterization of HC Clusters in *M. rosenbergii*

The hemocyte differentiation trajectory revealed that some granulocyte (GC) branches differentiated into hyalinocytes (HC). However, it remains unclear whether certain HC cells identified in this study are still in an intermediate state of differentiation and how their gene expression patterns change during this process. To investigate this possibility, we re-clustered the HC population (including clusters 6, 8, and 11) separately (Fig. 8A).

**Figure 8.**
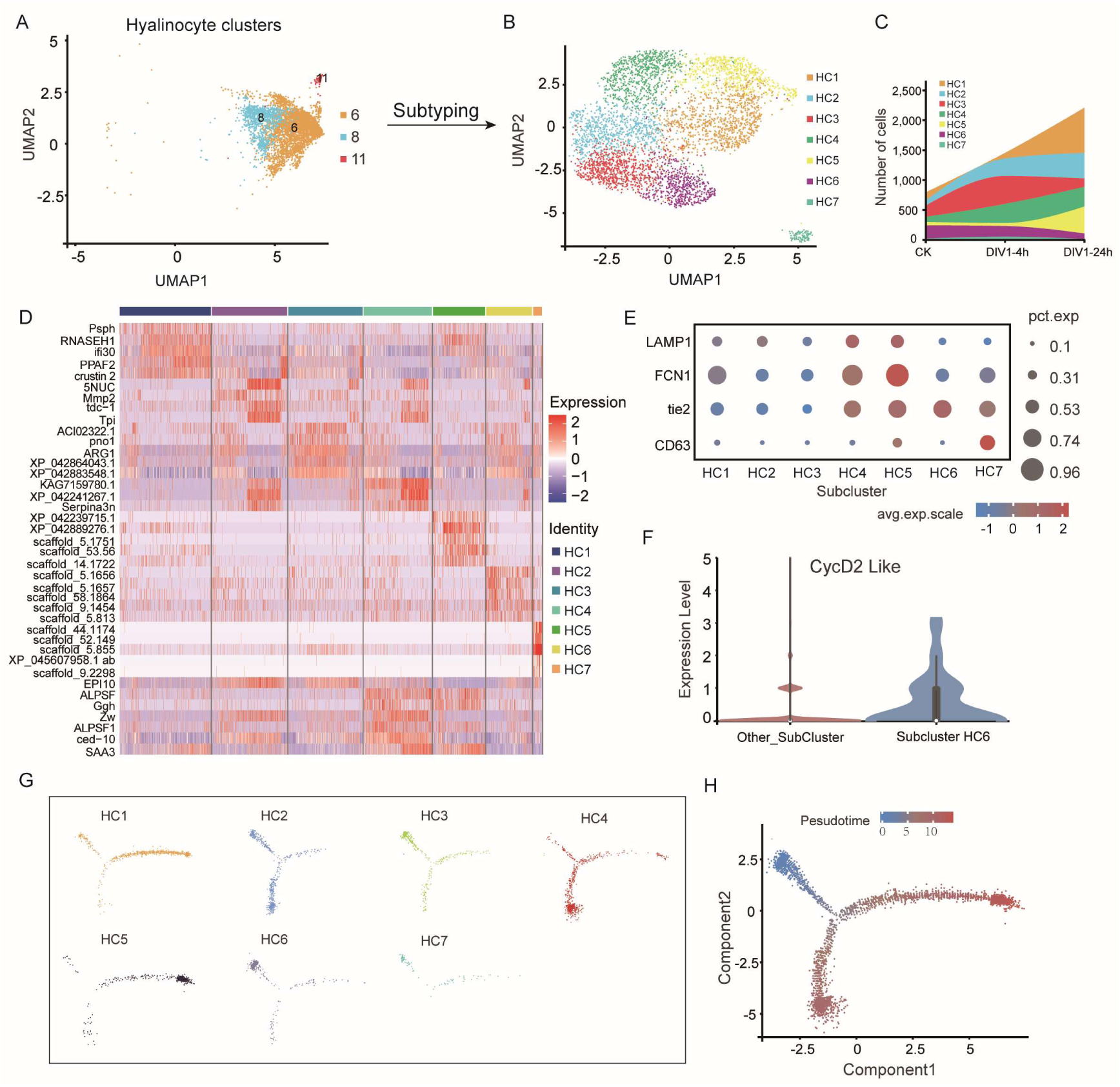
Subtyping of hyalinocyte subpopulation and characterization of their molecular profiles. (A) UMAP visualization of hyalinocyte subpopulation, identifying clusters 6, 8, and 11 for subtyping analysis. (B) UMAP plot showing the further subdivision of hyalinocyte subpopulation into seven subclusters (HC1–HC7). (C) Stacked area plot depicting the distribution of cells across clusters and time points (CK, DIV1-4h, and DIV1-24h). (D) Heat maps representing the gene expression profiles of subclusters (HC1–HC7), with selected marker genes labeled. (E) Dot plot showing the average expression levels and percentages of cells expressing key marker genes (LAMP1, FCN1, tie2, CD63) across the seven subclusters. (F) The expression levels of cell cycle-related gene *Cyclin D2* in subcluster HC5 compared to other subclusters by violin plot. (G) Pseudo-time trajectory analysis showing the progression of cells within the seven subclusters (HC1–HC7). (H) Mapping of pseudo-time value on differentiation trajectory of hyalinocyte subpopulation, as the color changes from blue to red, the value of the pseudo-time increases.

The re-clustering analysis identified seven subclusters, designated HC1 to HC7, among which HC1, HC2, HC4, and HC5 exhibited a continuous increase in cell numbers following DIV1 infection (Figs. 8B, 8C). Heatmap analysis of marker gene expression revealed that HC1 displayed the highest expression levels of the GC marker genes *PPAF2* and *Crustin2*. Additionally, genes associated with serine protease inhibition (*Serpina3n*), immune-related proteins (*EPI10*), nucleotide metabolism (*5’-nucleotidase-like*, *5NUC*), and energy production (*triosephosphate isomerase*, *Tpi*) were highly expressed in HC2 and HC4 (Fig. 8D).

Further analysis identified co-expressed genes in the HC4 and HC5 subclusters, including *ALF*, *γ-glutamate hydrolase-like gene* (*Ggh*), *LAMP1*, and phagocytosis-related genes (*FCN1* and *Tie2*). In contrast, these genes were expressed at relatively low levels in the HC2 subcluster (Figs. 8D, 8E). Moreover, cell cycle analysis showed that *CycD* expression was significantly higher in HC6 than in other subclusters (Fig. 8F), suggesting that HC6 represents an undifferentiated progenitor state. Based on this, we designated HC6 as the starting population and applied Monocle analysis to reconstruct differentiation trajectories for the seven subclusters.

The trajectory analysis revealed a transition from undifferentiated states (HC6 and HC2) toward two major differentiation branches (Fig. 8G). Branch 1 predominantly gave rise to HC1 and HC5, whereas Branch 2 mainly differentiated into HC4, which plays a key role in the peptide production and signaling transduction (Fig. 8H; Supplementary Figs. 7A, 7B). Additionally, we identified cells at the “junction” between branches, likely representing intermediate transitional states that complement HC4 (Fig. 8G).

Taken together, these findings demonstrate the transcriptional heterogeneity and dynamic differentiation of hyalinocytes during DIV1 infection, highlighting the complex interplay between viral infection and hemocyte lineage commitment.

## 3. Discussion

Decapod Iridescent Virus 1 (DIV1) is an emerging viral pathogen that primarily targets the hematopoietic tissues and hemocytes of crustaceans, leading to significant economic losses in aquaculture (1, 43). Understanding the specific hemocyte types targeted by DIV1 is essential for elucidating its pathogenesis and developing effective disease management strategies. While previous transcriptomic studies have provided insights into shrimp hemocyte immune responses, they lacked a comprehensive cellular and molecular perspective (7, 8). In this study, we applied single-cell RNA sequencing (scRNA-seq) to characterize 12 transcriptionally distinct hemocyte subtypes in *M. rosenbergii*, providing novel insights into their differentiation, immune functions, and interactions with DIV1. Our findings revealed significant shifts in hemocyte populations during DIV1 infection, with clusters 0, 5, 6, and 8 increasing in proportion. DIV1 primarily targeted clusters 0, 5, and 8, which were enriched in pathways associated with reactive oxygen species (ROS), infection response, and phagocytosis. The identification of functionally distinct hemocyte subpopulations, such as the antiviral HC4 cluster, and the discovery of resistant clusters (9 and 10), highlighting the complexity of hemocyte diversity and their differential roles in combating viral infections. These findings not only advance our understanding of shrimp immune responses but also provide a framework for disease resistance strategies in aquaculture.

### Hemocyte Diversity and Subtype-Specific Marker Genes

Hemocytes exhibit significant diversity in morphology and function, reflecting their specialized roles in various physiological processes (44, 45). Traditionally, shrimp hemocytes, similar to their counterparts in *Drosophila* and mosquitoes, have been classified into three main types: granulocytes (GC), semi-granulocytes (SGC), and hyalinocytes (HC) based on morphology (11, 12, 23, 46). However, advances in scRNA-seq have expanded this classification, enabling more precise identification of hemocyte subpopulations and their functional specialization. For example, scRNA-seq in *Drosophila* has revealed eight hemocyte subgroups, while studies in shrimp *L. vannamei* identified classic hemocyte types alongside novel subpopulations, including macrophage-like hemocytes (47, 48). Consistent with these findings, our study identified 12 transcriptionally distinct hemocyte subtypes in *M. rosenbergii* based on gene expression profiles. Prohemocytes (PH) in clusters 2, 7, and 9 were characterized by the expression of *ASPM*, *INCENP*, and *KIF20A*. Granulocytes (GC) in clusters 0 and 5 were defined by the expression of *PPAF*, *Crustin*, and *GPX3*. Semi-granulocytes (SGC) in clusters 3, 4, and 10 expressed *MUC2*, *SOD2*, and *Casp2*, while hyalinocytes (HC) in clusters 6 and 8 expressed *FCN1*, *Tie2*, and *CD63*. These results underscore the utility of scRNA-seq in resolving hemocyte diversity and their specialized immune functions. Furthermore, the identification of resistant hemocyte clusters (9 and 10) offers potential targets for genetic selection in shrimp breeding programs aimed at enhancing disease resistance.

### Hemocyte Differentiation Dynamics and Functional Adaptations During DIV1 Infection

Hemocyte differentiation is a dynamic process that is significantly influenced by viral infection (35, 49, 50). Our pseudo-temporal analysis revealed that DIV1 infection actively drives hemocyte differentiation, altering developmental trajectories and functional roles. We observed a rapid transition of prohemocytes (PH) and semi-granulocytes (SGC) into terminally differentiated granulocytes (GC) and hyalinocytes (HC), likely as part of an immune mobilization strategy against DIV1. Interestingly, transitional states were observed where GC cells differentiated into HC, reflecting the plasticity of hemocyte differentiation under viral stress. The recruitment and activation of terminally differentiated cells were accompanied by upregulation of immune-related pathways, reinforcing their adaptive roles in immune defense (51). Re-clustering of HC subpopulations revealed seven distinct subclusters (HC1–HC7), with HC4 emerging as a key antiviral population enriched in genes related to phagocytosis (*FCN1*, *Tie2*) (36, 52). These findings suggest that DIV1 infection accelerates hemocyte differentiation, reallocating immune resources to enhance host defenses. This knowledge has practical implications for disease control, as immunostimulatory treatments could be designed to enhance the recruitment of functional hemocytes during infection. Additionally, understanding how viral infections reshape hemocyte differentiation may inform selective breeding programs to improve shrimp resilience.

### Virus-Host Interactions: Viral Gene Expression and Hemocyte Susceptibility

Our scRNA-seq analysis revealed distinct viral targeting patterns across hemocyte clusters. DIV1 transcripts were detected in all hemocyte clusters except clusters 9 and 10, suggesting that these clusters possess intrinsic resistance mechanisms. In contrast, clusters 0, 5 and 8 exhibited the highest levels of viral gene expression, indicating that late-stage differentiated hemocytes are the most susceptible to infection. This preference for mature hemocytes may be attributed to their enhanced transcriptional and metabolic activity, which viruses can exploit for replication (35, 53, 54). Immediate-early (IE) genes, such as *ICP-46* (*73R*), were highly expressed in cluster 11, suggesting its involvement in the initiation of viral replication. Meanwhile, late-stage viral structural proteins, including *169R*, *16R*, and *148L*, were predominantly expressed in clusters 5 and 8. These proteins are crucial for viral assembly, highlighting the importance of these hemocyte subpopulations in sustaining viral propagation (41, 55, 56). The identification of resistant hemocyte clusters (9 and 10) presents an exciting opportunity for future research. Investigating the molecular basis of their resistance could lead to novel strategies for enhancing shrimp immunity, such as the selection of disease-resistant lines in breeding programs.

### Antiviral Responses and Immune Effector Genes

Clusters 0, 5, 6, and 8 were identified as key players in the antiviral response to DIV1 infection. These clusters exhibited significant enrichment of immune-related pathways, including antimicrobial peptides (AMPs), the prophenoloxidase (proPO) system, zinc finger proteins (ZNFs), and heat shock proteins (HSPs). AMPs act as a first line of defense by targeting various stages of the viral life cycle, while ZNFs mediate antiviral responses through nucleic acid binding and protein interactions (38, 57–59). HSPs, in addition to their chaperone functions, are actively involved in immune signaling (60–62). Key immune genes such as *PPO2*, *RAB11B*, *LAMP*, and *Dorsal* were significantly upregulated in these clusters, and functional knockdown experiments confirmed their roles in restricting viral replication and enhancing host survival. These findings highlight potential targets for immunostimulatory treatments or genetic improvement programs to boost shrimp resistance against viral infections.

In conclusion, this study provides the first comprehensive analysis of hemocyte differentiation and immune responses in *M. rosenbergii* during DIV1 infection. By leveraging scRNA-seq, we identified 12 transcriptionally distinct hemocyte subtypes, elucidated key functional pathways, and revealed the complex interplay between viral strategies and host immune defenses (Fig. 9). The insights into hemocyte subtypes, differentiation trajectories, and resistance mechanisms offer a foundation for improving shrimp breeding programs and developing targeted interventions for disease management in aquaculture.

**Figure 9.**
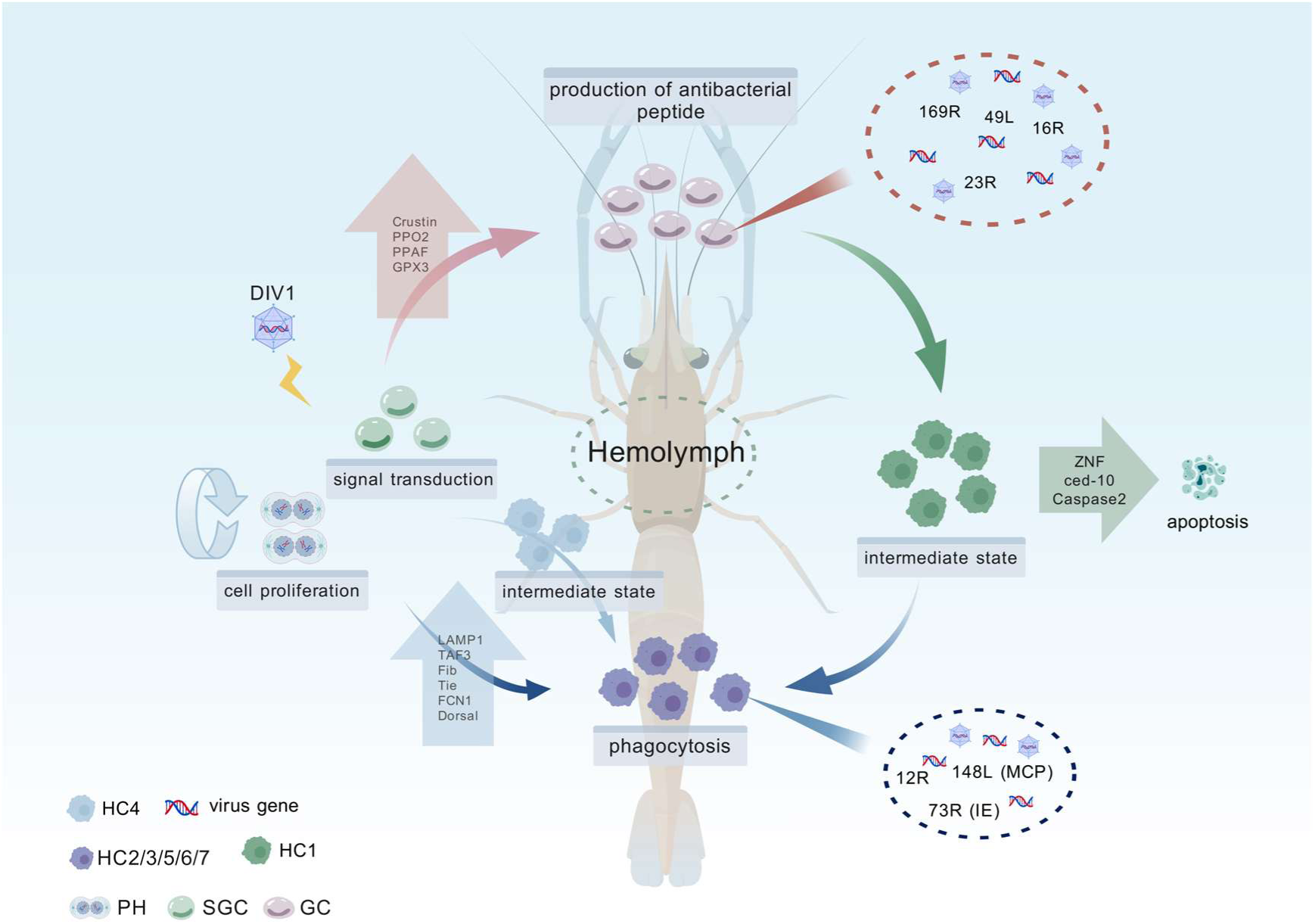
Schematic diagram of the functional differentiation of hemocytes during DIV1 infection in *M. rosenbergii*. Upon DIV1 infection, prohemocytes (PH) and semi-granulocytes (SGC) undergo differentiation into granulocytes (GC) and hyalinocytes (HC), leading to significant shifts in hemocyte composition. Throughout this process, intermediate transitional subpopulations, such as HC1 and HC4, emerge, co-expressing marker genes characteristic of both GC and SGC, respectively, indicating a dynamic differentiation state. During infection, PH cells contribute primarily to cell proliferation, while SGC cells play a key role in signal transduction. As infection progresses, the proportions of GC and HC cells steadily increase. Functionally, GC cells are responsible for antimicrobial peptides (AMPs) production, whereas HC cells are actively involved in phagocytosis. However, DIV1 selectively exploits these terminally differentiated hemocytes, utilizing GC and HC cells as replication hubs for viral gene expression and propagation. In the later stages of infection, the upregulation of apoptosis-related genes, including *ZNF*, *ced-10*, and *caspase-2*, triggers programmed cell death, which may serve as a host defense mechanism to limit viral spread. This figure is created with BioGDP.com (72).

## 4. Materials and methods

### 4.1 Animal and virus

The *M. rosenbergii*, approximately 25 ± 2.7 *g*, utilized in this experiment were procured from Zhuhai, Guangdong Province, China. The experimental shrimp were subjected to random sampling and testing to ascertain that they did not carry any pathogens, including DIV1, WSSV, and other pathogens according to the previous study (63). We cultured the shrimp in freshwater culture tanks equipped with oxygenation device, filtration system and kept at 26 °C for one week. During the experimental period, 10% of the water was changed daily. Shrimp were fed twice, and water body ammonia nitrogen and nitrite levels were tested regularly to bring them within safe thresholds.

The DIV1 (GenBank: ASM478855v1) used in this experiment was obtained by grinding muscle extraction from diseased shrimp infected with DIV1, and the DIV1 inoculum was prepared and quantified according to the previous method of our lab (14). A DIV1-infected challenging group and a control group (CK) were set up. 50 μL of DIV1 suspension diluted at a concentration of 1.02 × 10^7^ copies/μL was injected into the shrimp in challenging group through intra-muscular (IM) injection. 50 μL of PBS was injected into the shrimp in negative control group through IM injection. The DIV1 challenging group was subjected to hemolymph suspension preparation at 4 and 24 hpi, respectively.

### 4.2 Preparation of shrimp hemolymph in suspension

*M. rosenbergii* were randomly selected from various groups, then 2 mL of cold anticoagulant (27 mM Na citrate, 336 mM NaCl, 115 mM glucose, 9 mM EDTA, pH 7.2) and shrimp pericardial hemolymph were withdrawn using a 5mL syringe in a ratio of 1:1, respectively. They were mixed well and pooled in 15 mL centrifuge tubes, centrifuged at 600 *g* for 5 min at 4 °C, then the supernatant was discarded. The cells were resuspended using 3 mL of modified *M. rosenbergii* specific balanced salt solution (26.235 g NaCL, 0.745 g KCL, 2.84 g Na_2_HPO_4_, 3.09 g HEPES, 1.5 g BSA stirred, dissolved and fixed to 1000 mL, and sterilized by 0.22 μm filtration). After uniform blowing, the cells were filtered through a 40 μm cell sieve, and the filtrate was centrifuged at 600 *g* for 5 min at 4 °C to recover the cells. The supernatant was discarded and the cells were resuspended by adding 1 mL of pre-cooled modified *M. rosenbergii* specific balanced salt solution to the cell precipitate, and the cells were recovered by filtering the cells through a 40 μm cell sieve, then the filtrate was centrifuged at 600 *g* for 5 min at 4 °C. The supernatant was discarded, and the hemocytes were resuspended with pre-cooled modified *M. rosenbergii* special balanced salt solution. 10 μL of cell suspension was collected and stained with 0.4% Tepan blue for 1 min. The concentration and viability of the cells were observed using a cell counter plate under the microscope, and the concentration of the cells was diluted with *M. rosenbergii* special balanced salt solution to a concentration of 1500 ∼ 2000 cells/μL after the two parameters.

### 4.3 Single-cell encapsulation and sequencing

After mixing the hemocyte suspensions with 0.4% Taipan blue staining solution in a 5:1 ratio, the total number of hemocytes and cell viability were counted using a cell counter, and qualified samples with survival rate ≥ 90% were examined. The concentration of cells was adjusted to 1500 ∼ 2000 cells/μL and waited for loading on the machine. Single-cell isolation and library construction of hemocytes from PBS-treated and DIV1-infected groups of *M. rosenbergii* were performed using a 10 × Genomics Chromium Controller system (64). The main processes of single cell isolation include binding of gel beads to a mixture of cells and enzymes, single cell encapsulation of surfactant droplets, formation of water-in-oil Gel Beads In-Emulsions (GEMs), collection and permeabilization of the GEMs, PCR amplification of reverse transcribed cDNA, and construction of a standard library. Then, an indexed sequencing library of amplified cDNA from each cell in each sample was constructed separately using the reagents in the Chromium Single Cell 3’ Library v3 Kit.

Libraries were uploaded to the Illumina platform for high-throughput sequencing, and barcoded sequencing libraries of each sample after quantitative characterization were uploaded to the HiSeq2500 (Illumina, San Diego, CA), where each sample was individually sequenced and collected using custom double-ended sequencing.

### 4.4 Data filtering, normalization, and standard process analysis

Cell Ranger software was used for quality control as well as reference genome comparison, followed by validated barcode confirmation for distinguishing between different cells, and UMI for labeling unique transcription events. The following criteria were used to filter cells: (1) gene counts ˂ 550 or gene count ˃ 2600 per cell, and (2) UMI counts > 11,000 per cell.

After removing unwanted cells from the dataset, we employed a global-scaling normalization method “LogNormalize” that normalizes the gene expression measurements for each cell by the total expression, multiplies this by a scale factor (10,000 by default), and log-transforms the results. The formula is shown as follows:

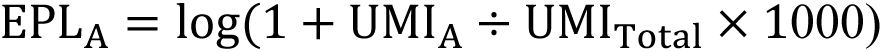

Among them, EPL_A_ is the normalized expression value of gene A in the target cell, UMI_A_ is the UMI number of gene A in the target cell, and UMI_Total_ is the sum of the effective UMI number in the target cell. It must be noted that the UMI_Total_ performed in the gene quantification does not include UMI_DIV1_. log is the natural logarithm of the base number e.

We used library size normalization methods to eliminate the influence of sequencing depth or cell size on expression between samples. Harmony algorithm was used to adjust and correct for batch effects due to experimental conditions, sequencing platforms, and processing times.

Standard process analysis of gene-cell expression matrices was performed using Seurat, including quality control and filtering of the data, normalization of expression data by the Normalize function, principal component analysis by RunPCA, cell clustering by the FindNeighbors function, and clustering and subclustering by the Louvain algorithm. The final cellular clustering results were visualized by the downscaled t-distributed Stochastic Neighbor Embedding (t-SNE) plot or Uniform Manifold Approximation and Projection (UMAP) plot.

### 4.5 Differentially expressed genes analysis per cluster

Differential expression genes (DEGs) of each cell subpopulation were screened and analyzed by Seurat’s Find Markers under the following conditions: (1) gene expression multiplicity |Log_2_ FC| ≥ 0.36, (2) *p* value ≤ 0.01, and (3) percentage of cells in a specific subpopulation where the gene is detected > 25% (65). The above threshold parameters are also applicable to the classification of DEGs between DIV1-infected group and control group. Furthermore, the DEGs for each cell cluster were evaluated using the Wilcoxon rank-sum test, and the resulting data were analyzed using GraphPad Prism (Version 9.0.0).

### 4.6 Data annotation and functional analysis

To identify various types of shrimp hemocytes, scRNA-seq data was automatically annotated using the Single R toolkit, while manual annotation was performed in combination with published literature and the Cell Marker database (http://117.50.127.228/CellMarker/index.html). The Gene Ontology (GO) and Kyoto Encyclopedia of Genes and Genomes (KEGG) enrichment analyses of the DEGs were conducted using the cluster profiler package’s enrich GO and KEGG functions, and the results were visualized using the Seurat package’s DotPlot and VlnPlot functions (66, 67).

### 4.7 Functional gene mining and differentiation trajectory construction

To classify key immune genes that may play a crucial role in antivirus process, STEM (Short Time-series Expression Miner, http://www.cs.cmu.edu/~jernst/stem) analysis was performed for the DEGs between different groups of cell clusters 0, 5, 6, and 8, respectively. To evaluate the differentiation relationship among various clusters, monocle 2 was used to construct differentiation trajectories for hemocyte clusters.

### 4.8 Viral genes analysis in cell clusters of the DIV1-infected group

To determine the state of DIV1 infection in each hemocyte cluster, we analyzed the expression of viral genes and viral load in the different cell clusters in the DIV1-infected group. Viral gene expression was analyzed using Cell Ranger, based on the DIV1 genome (GenBank: ASM478855v1). The “viral load” of a cell in scRNA-seq analysis was based on the number of UMIs that map to the DIV1 genome and expressed as a percentage of total UMI (UMI_DIV1_ + UMI_host_) content of a given cell (68). In this study, DIV1-infected shrimp hemocytes were divided into three categories of viral-load states: low (< 5%), medium (between 5 and 20%), and high (> 20%).

### 4.9 RNA interference (RNAi)

The T7 RiboMAX Express RNAi System kit (Promega, Shanghai, China) was used to generate double-stranded RNA (dsRNA) targeting the immune genes, and the dsGFP was used as a control. The specific primers used for dsRNA synthesis are listed in Supplementary Table 1. The integrity and quality of the synthesized dsRNA were verified using 1.5% agarose gel electrophoresis and quantified with a NanoDrop 2000 spectrophotometer (Thermo Scientific, Shanghai, China). Each *M. rosenbergii* received with 2 μg/g of body weight of the respective dsRNAs, dissolved in 50 μl of PBS. Control groups received injections of GFP dsRNA and PBS. Total RNA was extracted and assessed by quantitative PCR (qPCR) with corresponding primers for the evaluation of RNAi efficacy as described previously (69). For survival rate experiment, 48 hours after dsRNA injection, the shrimp were injected again with 1.02×10^7^ copies/μL of DIV1 or mock-challenged with PBS as a control. The survival rates of each group were recorded at 4-hour intervals. Differences in survival between groups were analyzed using the Mantel-Cox (log-rank χ^2^ test) method, employing GraphPad Prism software (GraphPad Software, La Jolla, CA, USA).

### 4.10 Quantitative PCR (qPCR)

qPCR assays were performed to assessed the mRNA levels of genes in RNAi experiments *in vivo*. The expression of *MrLAMP1* (GenBank accession no. PQ046648.1), *MrDorsal* (KX219631.1), *MrPPO2* (AY947400.1), and *MrRAB11B* (KP216761.1) were detected using LightCycler480 System (Roche, Basel, Germany) in a final reaction volume of 10 ml, which was comprised of 1 ml of 1:10 cDNA diluted with double-distilled H_2_O, 5 ml of 2 × Polarsignal SYBR Green qPCR mix (MIKX, MKG800), and 250 nM of specific primers (Supplemental Table 1). The cycling program were performed as described previously (70). The expression of each gene was calculated using the Livak (2^-ΔΔCT^) method after normalization to *Mrβ-Actin* (AF221096).

### 4.11 Detection of viral loads by absolute quantitative PCR

To measure viral titers in shrimp, absolute quantitative PCR (ab-qPCR) was used. Briefly, the hemocyte were collected from DIV1-infected shrimp. Hemocyte DNA was extracted with a Tissue DNA Isolation Mini Kit (Vazyme, Nanjing, China). The concentration of DIV1 genome copies was quantified by ab-qPCR, using the specific primers and the TaqMan fluorogenic probe, as listed in Supplementary Table 1 (71). To ensure accuracy, each shrimp sample underwent three replicates of ab-qPCR. The number of DIV1 genome copies was calculated and normalized against 0.1 μg of shrimp tissue DNA.

### 4.12 Western blotting

The hemocyte samples were lysed IP lysis buffer (Thermo, cat. no. 87787) with a protease and phosphatase inhibitor mixture. Samples were boiled for 10 min, separated on SDS-PAGE gels followed by transfer to polyvinylidene difluoride (PVDF) membranes. After blocking in 5% bovine serum albumin (BSA) in TBS with 0.1% Tween-20 (TBS-T). Then, the membranes were incubated with anti-MCP Abs, which produced by Gene Create (Wuhan, China), or anti-β-Actin Abs (Merck, KGaA, Darmstadt, Germany; cat. no. MAB1501). After washing in TBS-T, membranes were incubated for 1 h at RT with anti-rabbit IgG HRP-conjugate or anti-mouse IgG HRP-conjugate (Promega, Shanghai, China). Finally, the membranes were developed with the enhanced chemiluminescent (ECL) blotting substrate (Thermo Scientific, Shanghai, China), and the chemiluminescent signal was detected using the 5200 Chemiluminescence Imaging System (Tanon, Shanghai, China).

### 4.13 Histological analysis

Histological analysis was performed to detect the impact of the immune genes knockdown with DIV1 infection on tissue morphology. The hepatopancreas were collected and fixed using Davidson’s AFA fixative (30% ethanol, 22% formalin, and 11.5% acetic acid) at 24 h after dsRNA application. After fixing for 24 h, the tissues were dehydrated, embedded in paraffin, sectioned at 7 μm thickness, and stained using hematoxylin and eosin (H&E) as described previously (62). The slides were observed and images were captured under a Flexible, brightfield, fluorescent and fish scanner (Leica, Wetzlar, Germany).

### 4.14 Statistical Analyses

All data were presented as means ± SD. Student’s *t* test was used to calculate the comparisons between groups of numerical data at \**p* < 0.05, and \*\**p* < 0.01. For survival rates, data were subjected to statistical analysis using GraphPad Prism software (Graphpad, San Diego, CA, USA) to generate the Kaplan–Meier plot (log-rank χ^2^ test).

## Author contributions

Chaozheng Li, Jianguo He and Bang Xiao conceived and designed the experiments. Bang Xiao, Fang Kang, Taolin Fan, Qianqian Li, Manman Liu, Junming Pan, and Chaozheng Li performed the experiments and analyzed data. Bang Xiao, Fang Kang and Chaozheng Li wrote the draft manuscript. Bang Xiao, Chaozheng Li and Jianguo He acquired finding. Chaozheng Li was responsible for forming the hypothesis, project development, data coordination, and writing, finalizing, and submitting the manuscript. All authors discussed the results and approved the final version.

## Acknowledgement

This research was supported by National Natural Science Foundation of China (32373158/32441085/31930113), the open competition program of top ten critical priorities of Agricultural Science and Technology Innovation for the 14th Five-Year Plan of Guangdong Province (2022SDZG01), the Guangdong S&T programme (2024B1212060001), China Postdoctoral Science Foundation (2023M743987), and Guangdong Basic and Applied Basic Research Foundation (2023A1515110528). The funders had no role in study design, data collection and analysis, decision to publish, or preparation of the manuscript.

## Data availability

All reagents and experimental data are available within transparent methods or from corresponding author upon reasonable request.

## Ethics statement

All animal experiments were approved by Institutional Animal Care and Use Committee of Sun Yat-Sen University (Approval No. SYSU-IACUC-2024-B0640, 6 March 2024).

## Supplementary Figure

**Supplementary Figure 1.**
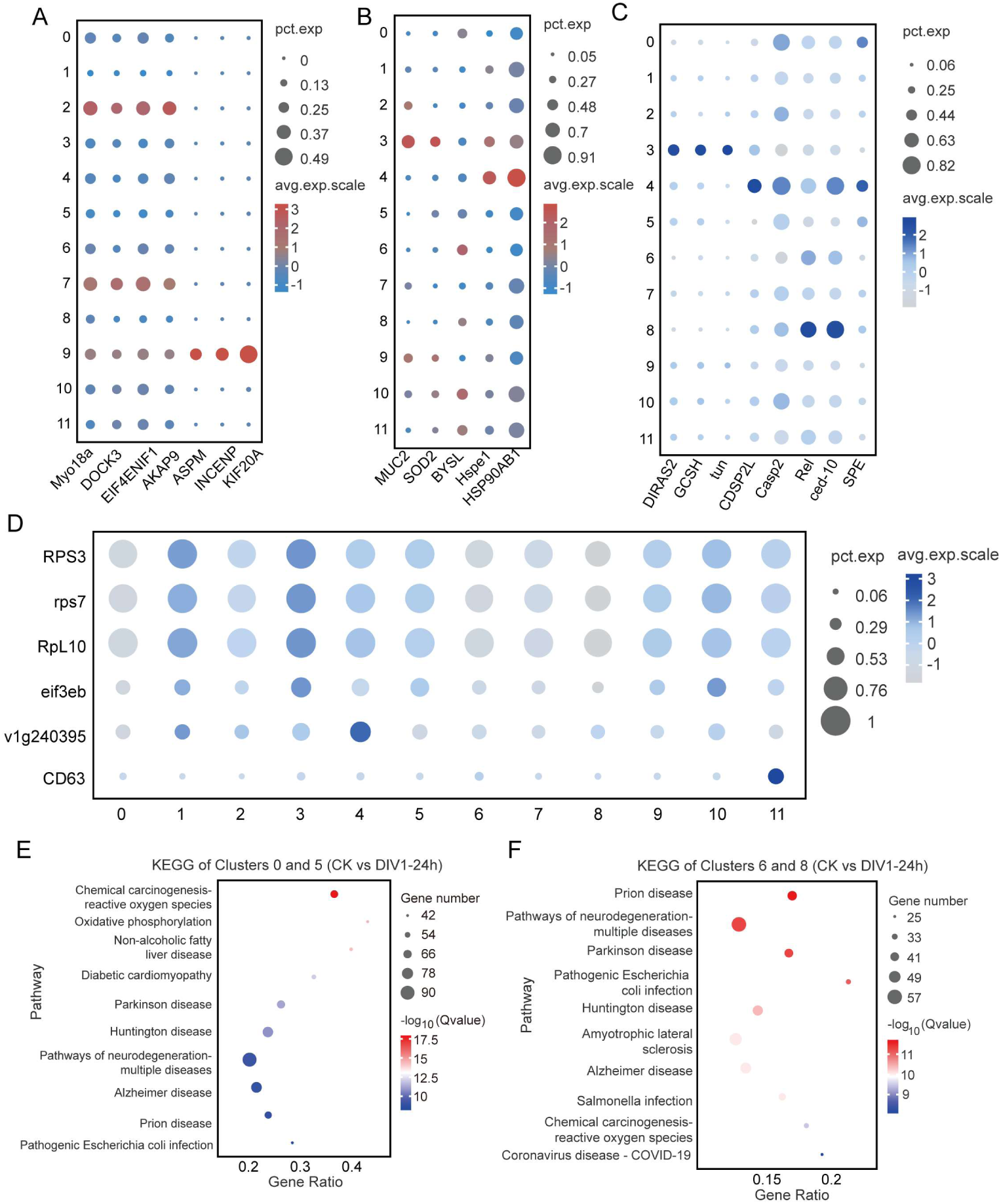
Genes especially high expressed in clusters 1, 2, 3, 4, 7, 9, 10, 11 and KEGG analysis of DEGs between group CK and DIV1-24h in clusters 0, 5 and 6, 8. (A) The high expression distribution of genes related to cell proliferation, division, and germline development in clusters 2, 7, and 9. (B) DEGs related to cytoskeletal formation, immune stress and cell development in cluster 3,4 and 10. (C) DEGs with SGC characterization in cluster 3, 4. (D) Expression profiles of key DEGs related to protein synthesis (*Rps3, rps7, RpL10, eif3eb, v1g240395*) and phagocytosis (*CD63*). (E, F) TOP 10 KEGG pathway enrichment analysis of differential genes in clusters 0 and 5 (E), clusters 6 and 8 (F) compared to the control group at 24 hpi. Dot size represents the number of genes involved, and color intensity indicates statistical significance.

**Supplementary Figure 2.**
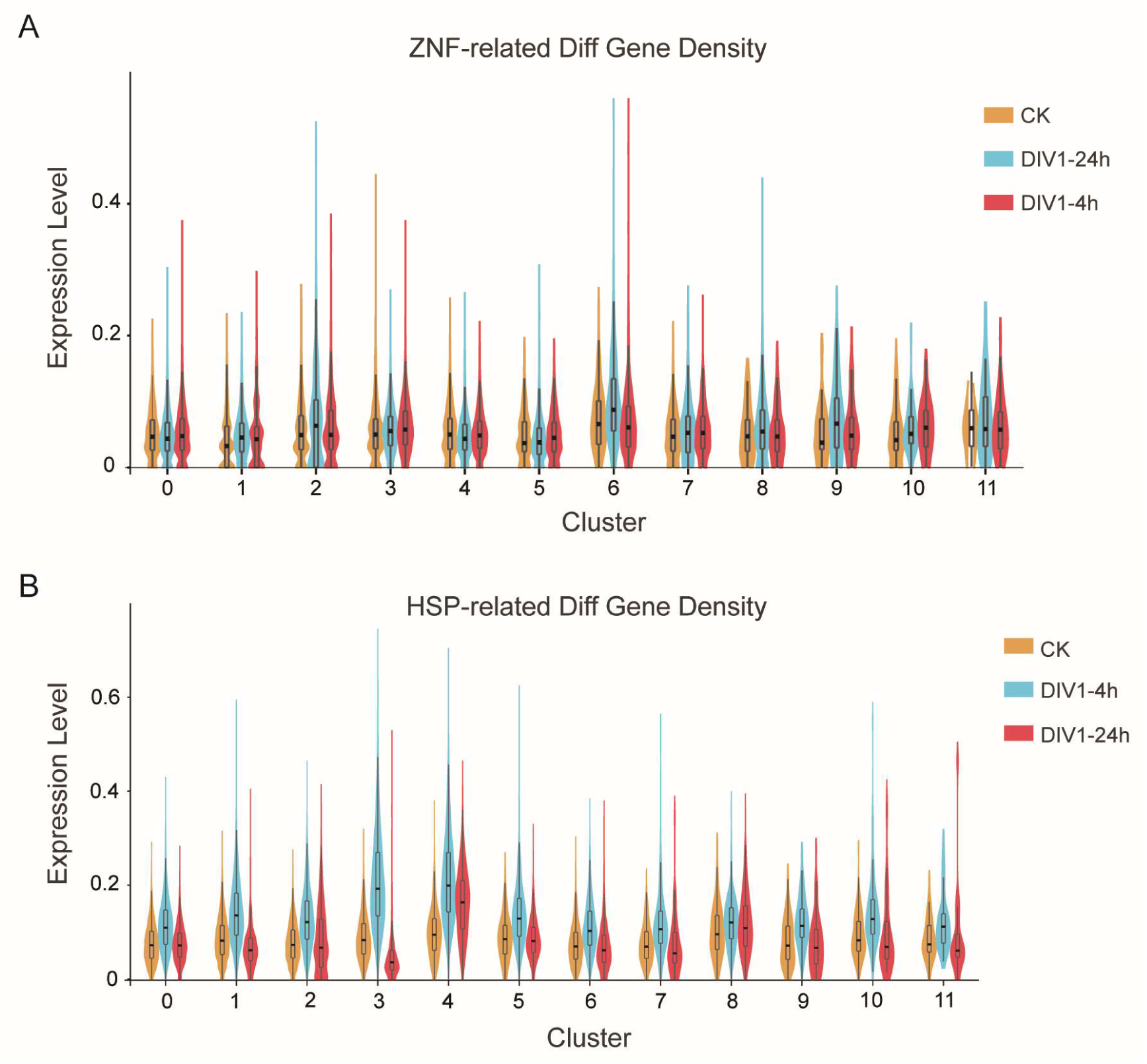
Expression profile of DEGs related to heat stress response or gene regulation between control group and DIV1-infection groups in each cluster. (A) The distribution of differentially expressed zinc finger proteins (ZNFs) in each cluster in different groups. (B) The distribution of differentially expressed heat shock proteins (HSPs) in each cluster in different groups.

**Supplementary Figure 3.**
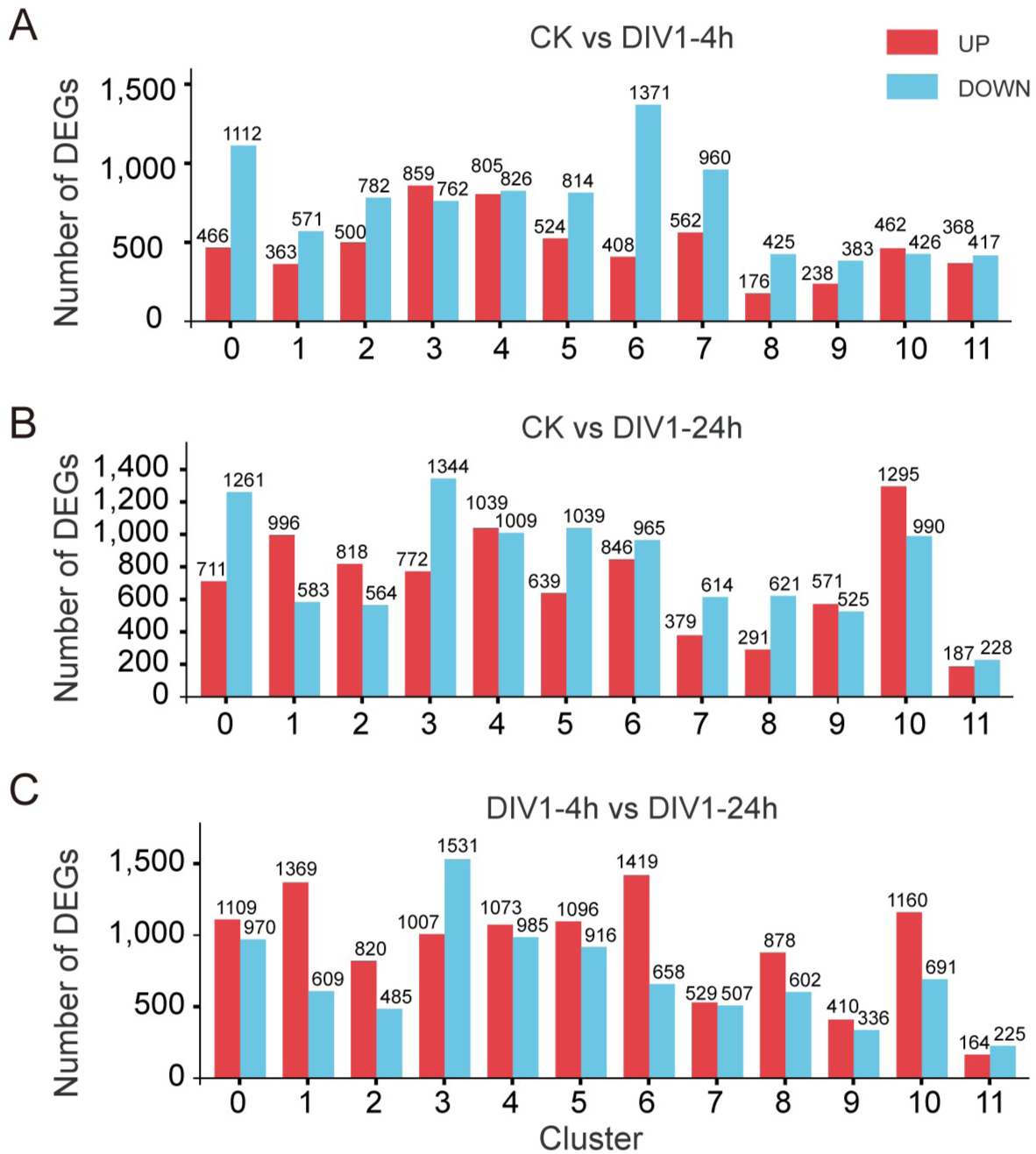
Overview of DEGs between DIV1-infected and control cells within shrimp hemocyte clusters. (A, B) Histogram showing all of the upregulated (red) and downregulated DEGs (blue) in DIV1-infected cells compared with control cells within all clusters at 4 (A) and 24 hpi (B). (C) Histogram showing all of the upregulated (red) and downregulated DEGs (blue) in group DIV1-24h cells compared with DIV1-4h cells.

**Supplementary Figure 4.**
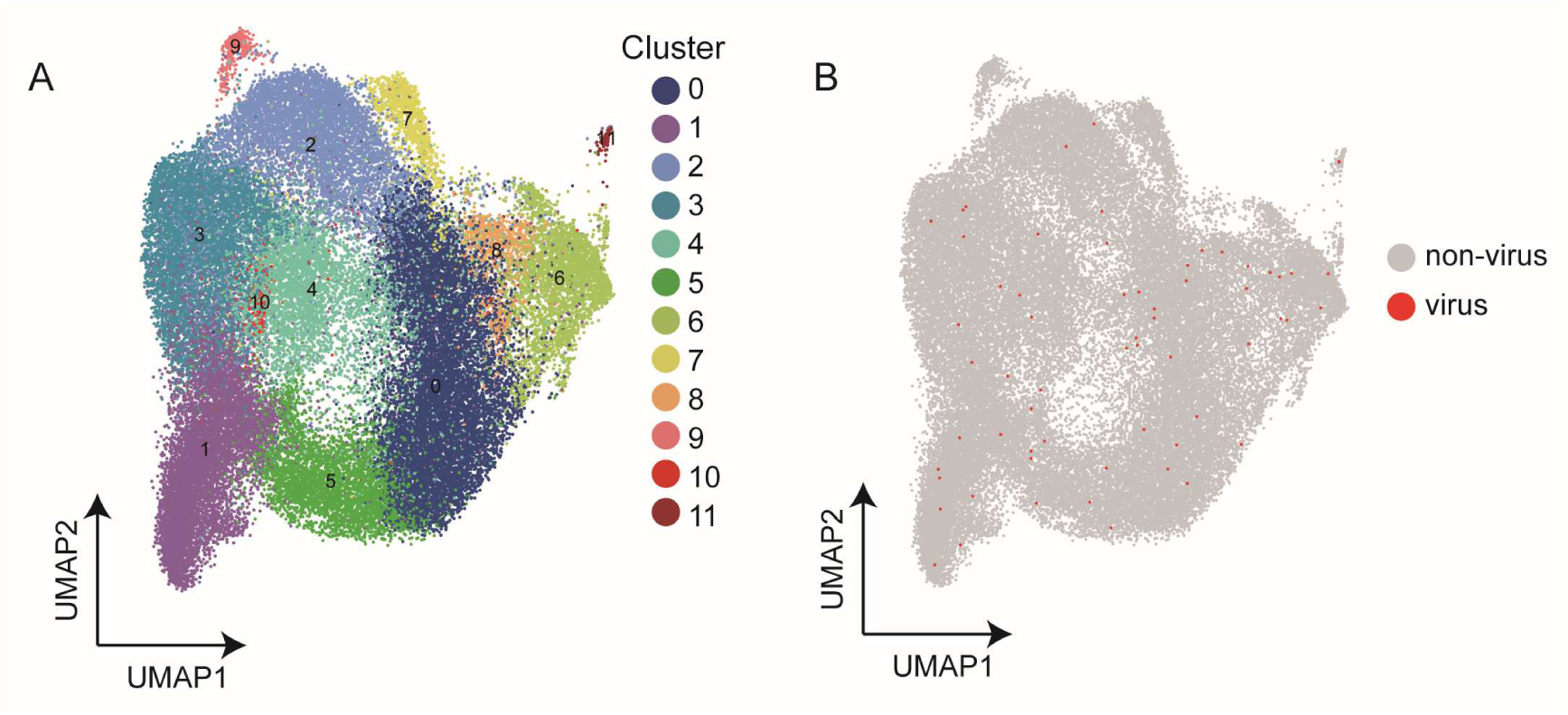
Detection of DIV1 genes of hemocyte cells within all clusters in DIV1-infected groups. (A) UMAP projection representing the 12 cell clusters identified in the shrimp hemocytes (unified set of control and DIV1 infection samples). (B) UMAP plot showing the expression atlas of all cells detected with virus genes in DIV1-infected samples, the dots in red represent the target cells.

**Supplementary Figure 5.**
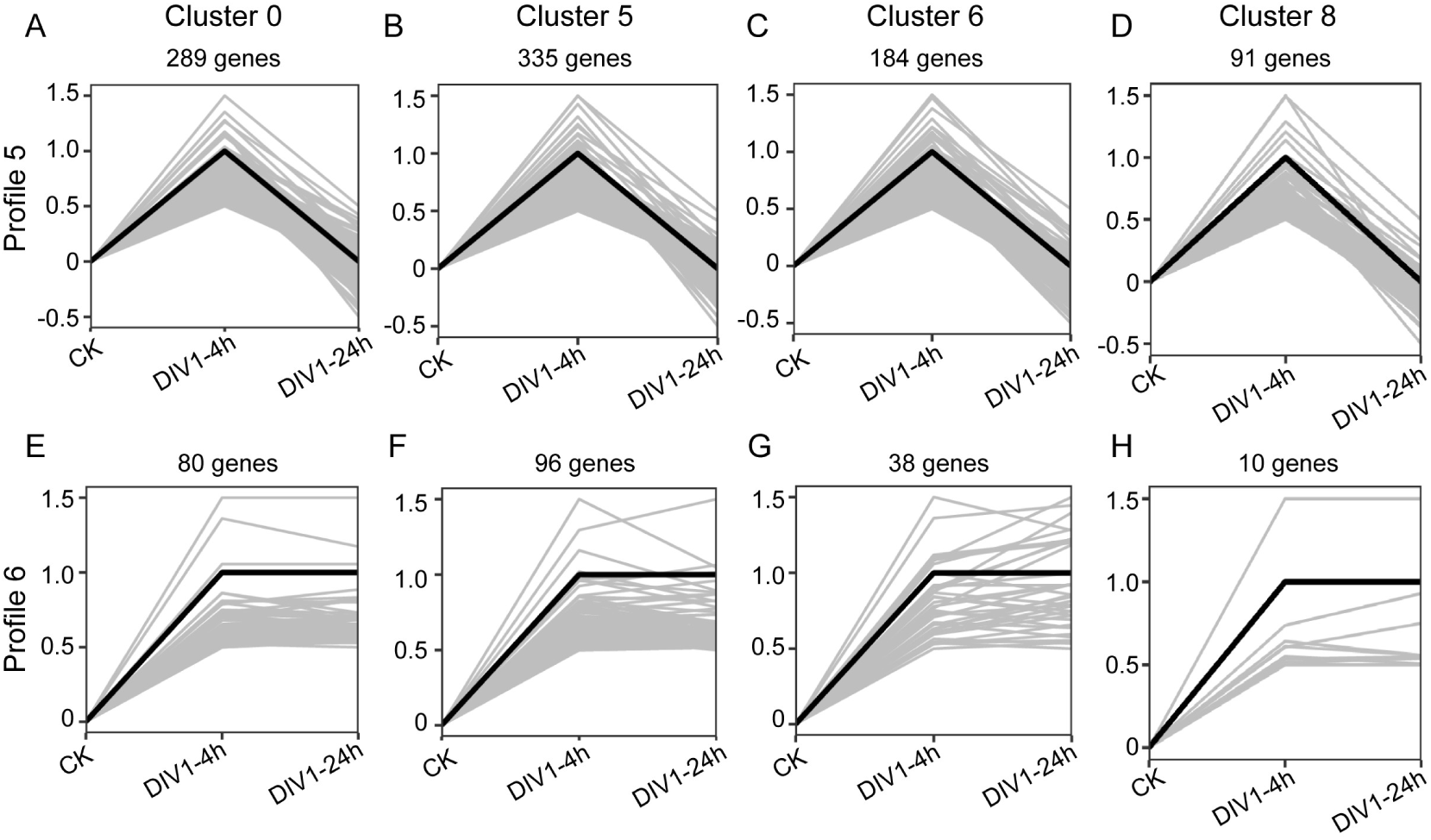
Patterns of gene expressions across three groups of hemocytes inferred by STEM analysis. (A-D) Profile 5 represented genes, in cluster 0 (A), cluster 5 (B), cluster 6 (C) and cluster 8 (D), which were highly up-regulated in group DIV1-4h. In each profile, the light gray lines represented the expression pattern of each gene, while the black line represented the expression tendency of all the genes. The number of genes belonging to each pattern was labeled above the profile. (E-H) Profile 6 represented genes, in cluster 0 (E), cluster 5 (F), cluster 6 (G) and cluster 8 (H), respectively, that were highly up-regulated in both groups DIV1-4h and DIV1-24h.

**Supplementary Figure 6.**
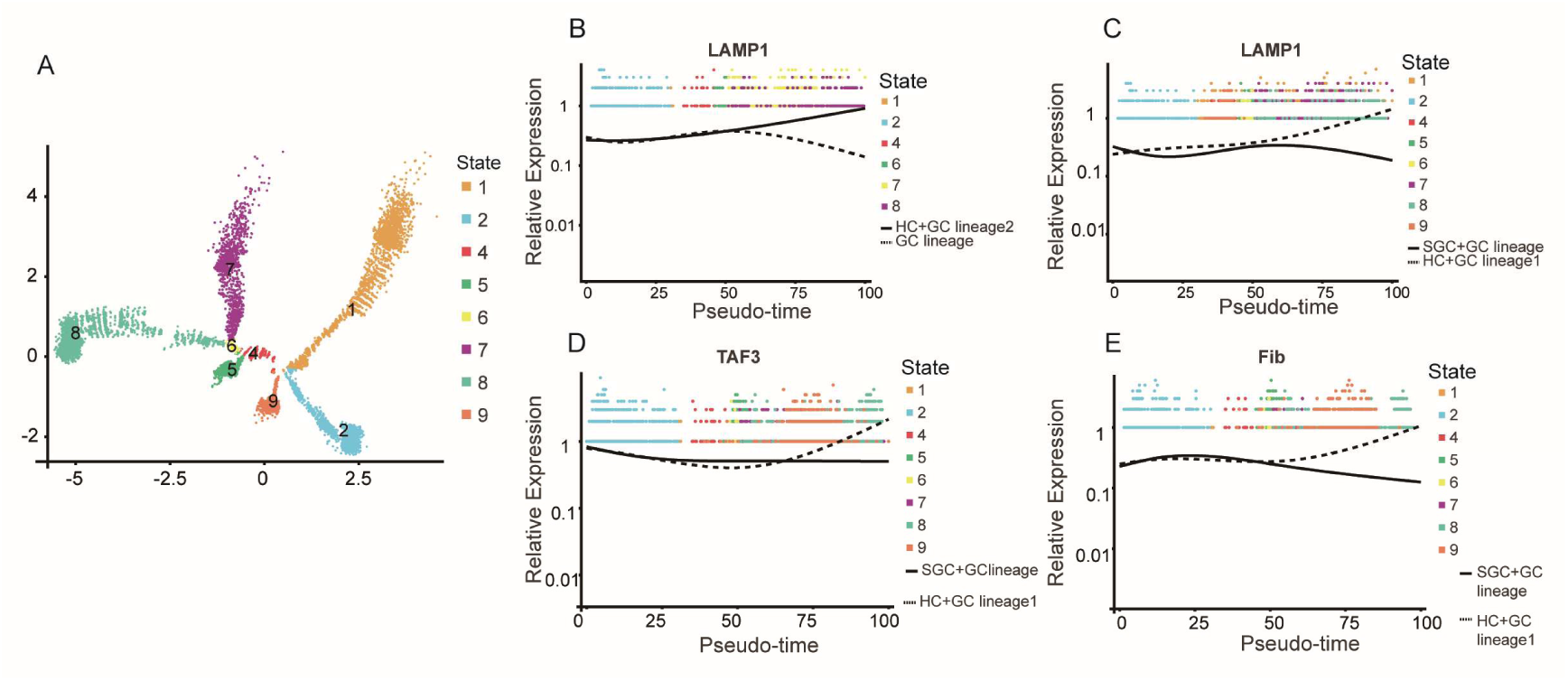
Trajectory distribution of genes of *LAMP1*, *TAF3*, and *Fib*. (A) Trajectory distribution of different differentiation states. (B, C) The expression trend of gene *LAMP1* in differentiation to HC branch 2 (B) or branch 1 (C) and other branches. (D, E) The expression trend of transcription factor *TAF3* (D)、*Fib* (E) in differentiation to HC branch and other branches.

**Supplementary Figure 7.**
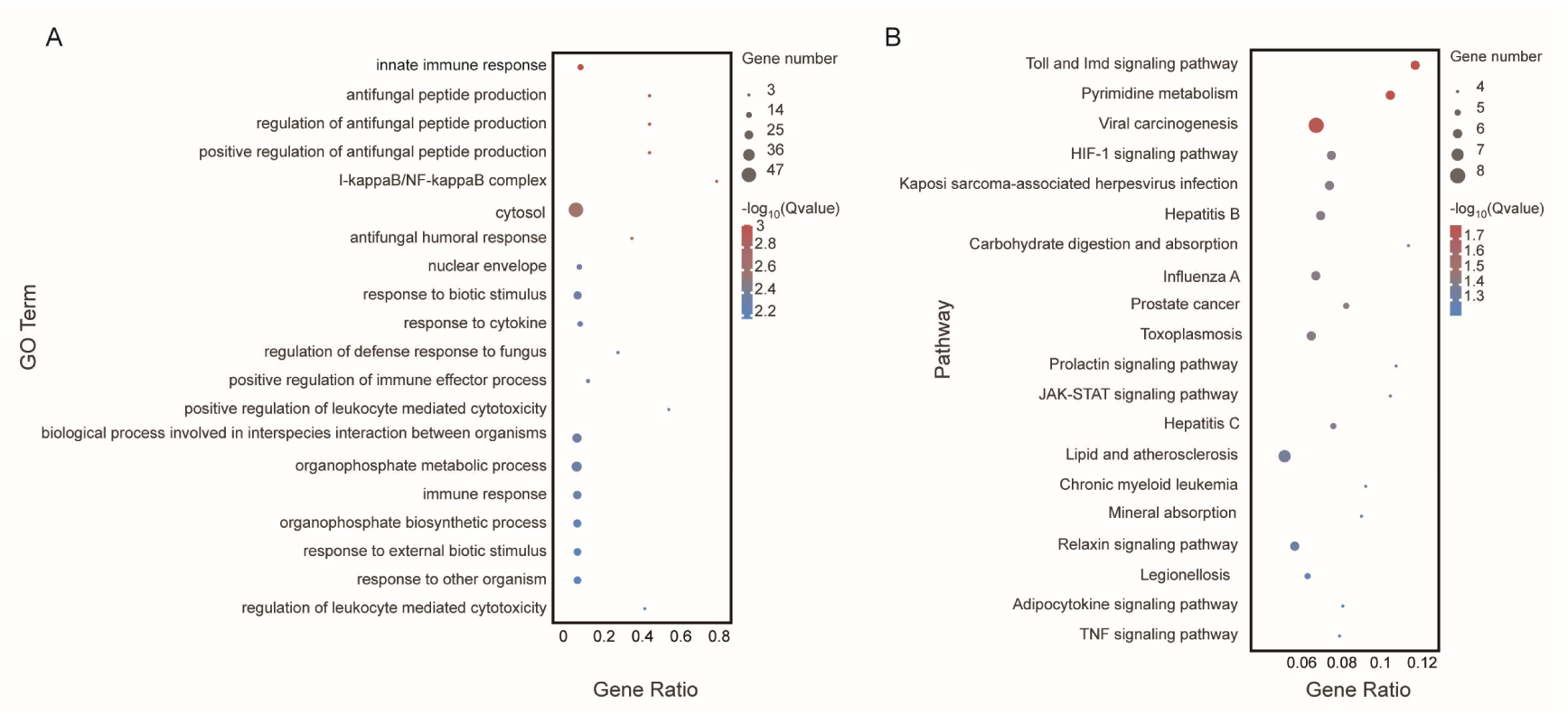
Function annotation of differential fate genes in branch 2 after HC re-clustering. (A, B) GO (A) and KEGG (B) enrichment analysis of differential fate genes in branch 2 after regrouping of hyalinocyte clusters.

